# Mapping the Organization and Morphology of Calcitonin Gene-Related Peptide (CGRP)-IR Axons in the Whole Mouse Stomach

**DOI:** 10.1101/2023.05.23.541811

**Authors:** Jichao Ma, Duyen Nguyen, Jazune Madas, Ariege Bizanti, Anas Mistareehi, Andrew M. Kwiat, Jin Chen, Mabelle Lin, Richard Christie, Peter Hunter, Maci Heal, Shane Baldwin, Susan Tappan, John B. Furness, Terry L. Powley, Zixi (Jack) Cheng

## Abstract

Nociceptive afferent axons innervate the stomach and send signals to the brain and spinal cord. Peripheral nociceptive afferents can be detected with a variety of markers [e.g., substance P (SP) and calcitonin gene-related peptide (CGRP)]. We recently examined the topographical organization and morphology of SP-immunoreactive (SP-IR) axons in the whole mouse stomach muscular layer. However, the distribution and morphological structure of CGRP-IR axons remain unclear. We used immunohistochemistry labeling and applied a combination of imaging techniques, including confocal and Zeiss Imager M2 microscopy, Neurolucida 360 tracing, and integration of axon tracing data into a 3D stomach scaffold to characterize CGRP-IR axons and terminals in the whole mouse stomach muscular layers. We found that: 1) CGRP-IR axons formed extensive terminal networks in both ventral and dorsal stomachs. 2) CGRP-IR axons densely innervated the blood vessels. 3) CGRP-IR axons ran in parallel with the longitudinal and circular muscles. Some axons ran at angles through the muscular layers. 4) They also formed varicose terminal contacts with individual myenteric ganglion neurons. 5) CGRP-IR occurred in DiI-labeled gastric-projecting neurons in the dorsal root and vagal nodose ganglia, indicating CGRP-IR axons were visceral afferent axons. 6) CGRP-IR axons did not colocalize with tyrosine hydroxylase (TH) or vesicular acetylcholine transporter (VAChT) axons in the stomach, indicating CGRP-IR axons were not visceral efferent axons. 7) CGRP-IR axons were traced and integrated into a 3D stomach scaffold. For the first time, we provided a topographical distribution map of CGRP-IR axon innervation of the whole stomach muscular layers at the cellular/axonal/varicosity scale.

## 1. Introduction

Abdominal pain is often related to aberrant neural control of the gastrointestinal (GI) tract. However, the anatomical organization and physiological processes of visceral nociceptive afferent axons have not been well elucidated. GI nociception is mainly mediated by sensory neurons in the spinal dorsal root ganglia (DRG) and, to a lesser extent, by the vagal nodose ganglia (VNG) (Frias and Merighi 2016, Gibbins et al. 1985, Sternini et al. 1992). In contrast to the extensive documentation of vagal afferent innervation of the stomach along with their mechanical receptors (intraganglionic laminar endings – IGLEs and intramuscular arrays – IMAs) in the rat and mouse (Fox et al. 2000, Furness et al. 2020, Powley et al. 2019, Powley 2021, Wang and Powley 2000), the topographical nociceptive afferent innervation in the stomach has not been well characterized. Substance P (SP), calcitonin gene-related peptide (CGRP), and transient receptor potential cation channel subfamily V member 1 (TRPV1) have been commonly used as three major nociceptive makers. Recently, we provided a comprehensive map of the SP-IR axon innervation in flat mounts of the whole mouse stomach muscular layers (Ma et al. 2023). Earlier studies (Ekblad et al. 1985, Furness et al. 1991, Green and Dockray 1988) showed that many SP-IR nerve fibers are of intrinsic origin, and our study supported this (Ma et al. 2023). Using the same methodology for the identification of morphology and distribution of nociceptive nerves, we applied immunohistochemical (IHC) labeling of CGRP to determine the distribution and morphology of CGRP-IR axon innervation in the whole stomach muscular layers of mice, as well as to determine the origin(s) of these axons.

CGRP, a peptide with 37 amino acid residues, is a neurotransmitter in capsaicin-sensitive afferent axons (Li et al. 2014, Luo et al. 2013, Russell et al. 2014) that extensively innervate the body and it has a dual role in sensory (nociceptive) and efferent (effector) function (Maggi 1995, Holzer 1998, Rosenfeld et al. 1983). CGRP and its receptors are widely expressed in both peripheral and central pain pathways (van Rossum et al. 1997). By the interaction with local gastric receptors, CGRP also exerts its physiological functions such as positive inotropic actions and vasodilation (Bell et al. 2016, Brain et al. 2004). In the GI tract, CGRP participates in the regulation of gastric blood flow, specifically as a strong vasodilator (Holzer and Guth 1991, Kee et al. 2018, Peskar et al. 2002). In addition to its vasodilation function, CGRP can induce muscle relaxation in rodent and human gastric smooth muscle (Katsoulis et al. 1989, Lee et al. 2020). Furthermore, the release of CGRP in the GI tract activates protective mechanisms that are important for a number of GI diseases including gastritis, gastroenteritis, and gastroparesis (Eysselein et al. 1992, Godlewski et al. 2010, Luo et al. 2013).

CGRP-IR fibers in the GI tract were observed in different species such as mouse, rat, guinea pig, hamster, axolotl, python, cat, dog, lamb, pig, and human using sectioned tissues (Chiocchetti et al. 2005, Dömötör et al. 2005, Furness et al. 2004, Gibbins et al. 1985, Hayakawa et al. 2009, Maake et al. 1999, Palus et al. 2018, Parkman et al. 1989, Poonyachoti et al. 2002, Rodrigo et al. 1985, Shochina et al. 1997, Sharrad et al. 2015, Sternini et al. 1992). Since sectioning of the tissues disrupted the continuity of the neural network of axons, some of these studies could not provide a complete view of the distribution and morphology of CGRP-IR axons and terminals in different regions and layers of the whole stomach. Given that a comprehensive view of the innervation pattern of CGRP-IR axons and terminals is important for the understanding of nociceptive processes, we used flat-mount preparations of the whole stomach muscle layers which retained the complete axon network. In addition to the holistic view, we also focused on specific gastric targets such as myenteric ganglia, longitudinal and circular muscles and blood vessels in different regions (Ma et al. 2023, Sharrad et al. 2015). Previously, we have studied the distribution pattern and morphology of CGRP-IR and SP-IR axons in flat mounts of the atria of the heart as well as SP-IR axons in flat mounts of the whole stomach muscular layers (Li et al. 2014, Ma et al. 2023). In this present study, we applied the same technique to determine the topographical distribution and morphology of CGRP-IR axons and terminals in flat mounts of the whole mouse stomach muscular layers. The determination of CGRP-IR axon morphology and distribution of the mouse stomach will add to our current understanding of this nociceptive marker and its function in the GI tract.

## 2. Materials and Method

### 2.1 Animals

Male C57BL/6J mice, n=19 with an age range from 12 to 16 weeks (RRID: IMSR_JAX000664, The Jackson Laboratory, Bar Harbor, ME) were used. Animals were kept in the animal room in which the dark/light cycle was set to 12/12 hours and water and food were supplied ad libitum. All procedures were carried out under the ethical guidelines of the University of Central Florida and approved by the Animal Care and Use Committee of the University of Central Florida.

### 2.2 Tissue preparation

Animals were anesthetized using isoflurane (4%, #029405, Covetrus North America, Dublin, OH). Sufficient depth of anesthesia was determined via the absence of the hind paw pinch withdrawal reflex. The abdomen and chest cavity were then opened with minimal incisions, and heparin (0.2 ml; 1,000 units/ml) was injected into the heart. Mice were then perfused with at least 150 mL 40°C phosphate-buffered saline (0.1 M PBS, pH = 7.4) via a blunt needle inserted into the left ventricle, and blood was drained by cutting the inferior vena cava. The perfusion solution was switched to 150 mL of ice-cold Zamboni’s fixative (15% of a saturated solution of picric acid, 2% paraformaldehyde in PBS, pH = 7.4) to fix the tissue.

After the perfusion, the stomach was removed from the visceral cavity and trimmed to include the distal lower esophageal sphincter and the proximal pylorus. The entire stomach was then opened with a longitudinal cut along the lesser and greater curvature into two equal halves (dorsal and ventral parts) and any food or residue in the stomach was removed by rinsing thoroughly with distilled water. The stomach was post-fixed in the same fixative for 12 hours. After post-fixation, each part of the stomach was cleaned, and the mucosal and submucosal layers were dissected from the muscular wall of the stomach. The myenteric plexus together with the longitudinal muscle layer (facing the serosa of the stomach) and circular muscle layer (facing the submucosal layer) comprised the stomach muscular wall.

The stomachs were supplied by the gastric and gastroepiploic artery branches. Whereas gastroepiploic branches supplied the stomach along the margin of the greater curvature, gastric branches supplied the central parts of the ventral and dorsal stomach. For simplicity, we removed most of the vasculature along the margin of the greater curvature to only show the gastric arteries (Tissue preparation in **Figure 1**). The left and right gastric arteries entered the ventral and dorsal stomach near the esophagus and pylorus. In the cardiac region where thick adipose tissues covered the area, these arteries penetrated through the muscular layer from the outer layer (the serosa), went into the submucosal layer, and then ramified and distributed in and between the muscular and submucosal layers. Together with the thick adipose tissues, the main branches of gastric arteries were removed, and only the branches attached to the immediate serosa remained.

**Figure 1.**
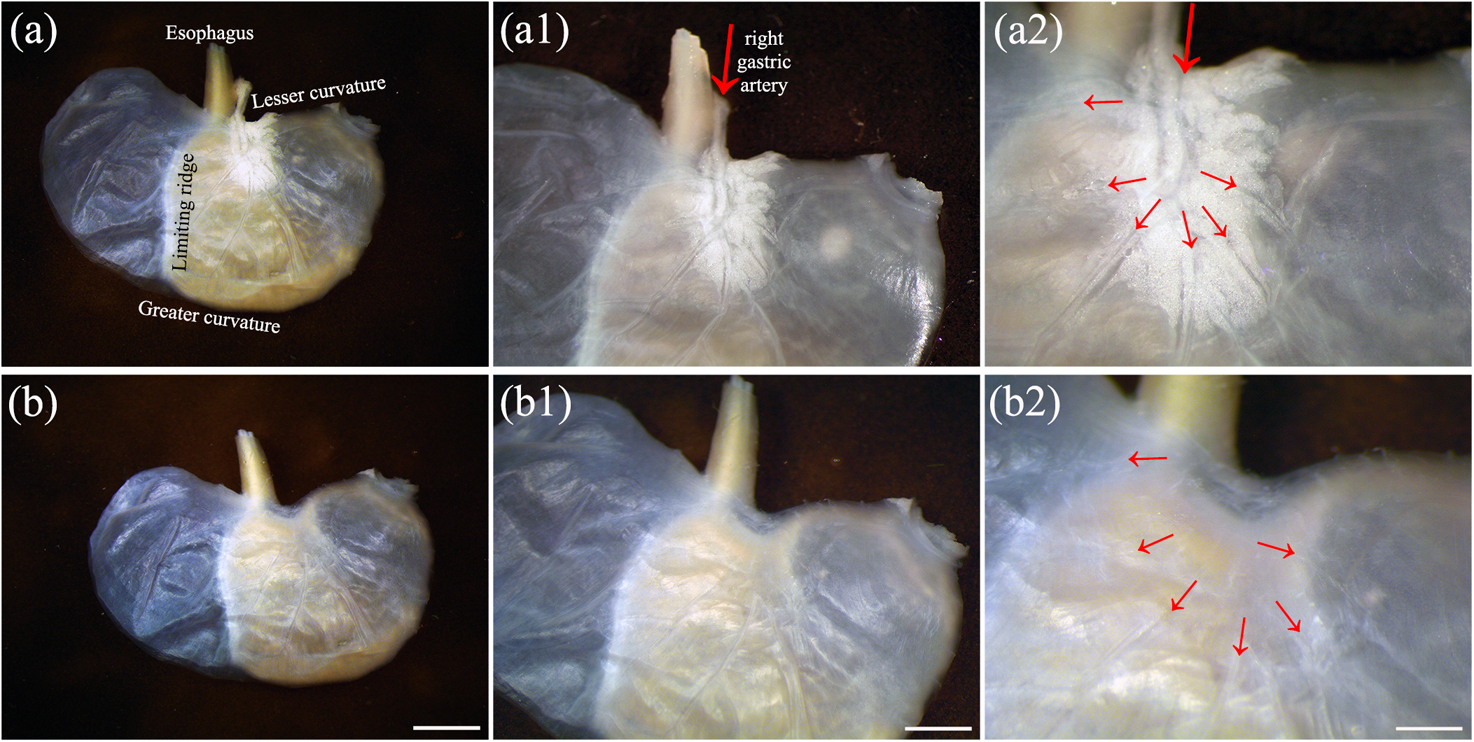
A flat-mount preparation of a representative stomach prior to immunohistochemistry. The stomach was cut longitudinally along the lesser and greater curvatures to yield two equal halves (dorsal and ventral). (a) A dorsal stomach with the serosa facing up and mucosa facing down, higher magnification in (a1) and (a2). The main branch of the right gastric artery (large red arrow), the connective tissues, and white adipose tissues at the entrance of the lower esophagus to the stomach are clearly seen. The small red arrows indicated the branches from the right gastric artery. (b) The main branch of the right gastric artery, the connective tissues, and adipose tissues at the entrance of the lower esophagus to the stomach were removed, only the vessel branches attached to the immediate serosa remained (small red arrows indicated the beginning of the remaining branches of the blood vessels). This stomach could be seen at higher magnification in (b1) and (b2). The stomach in (b) was used for immunohistochemistry. Scale bar in (b) = 4 mm, in (b1) = 2 mm, in (b2) = 1 mm. A higher resolution version of this figure is available in the Supporting Information.

### 2.3 Immunohistochemistry (IHC)

All IHC steps were performed using a 24-well plate on a shaker at room temperature (∼24 °C) in a dark environment. The IHC steps included washes, blocking, primary antibody, secondary antibody, and were carried out in different wells.

#### Single labeling immunofluorescence IHC

Following previously established protocols for whole mount processing (Li et al. 2014, Ma et al. 2023, Rysevaite et al. 2011), samples were washed 6x5 minutes in 0.1 M PBS (pH = 7.4) and immersed in blocking solution (0.1 M PBS containing 2% bovine serum albumin, 10% normal donkey serum, 2% Triton X-100, and 0.08% sodium azide) for 5 days. After that, the stomach was incubated in primary antibody monoclonal mouse anti-CGRP (ventral n=3, dorsal n=4; 1:150) or rabbit monoclonal anti-CGRP (ventral n=2, dorsal n=2; 1:150) for 4-5 days in a 0.1 M PBS solution containing 2% bovine serum albumin, 4% normal donkey serum, 0.5% Triton X-100, and 0.08% sodium azide. The tissue was extensively washed 6x5 minutes in 0.1 M PBS containing 0.5% Triton X-100 (PBS-T) to remove any unbound primary antibody, followed by incubation in fluorescent-labeled secondary antibody, donkey anti-mouse Alexa Fluor 488 (1:90) or donkey anti-rabbit Alexa Fluor 488 (1:90), respectively, for 3 days in 0.5% PBS-T. The tissue was washed 6x5 minutes in 0.1 M PBS and mounted on a slide with the serosal side facing up.

The tissue was flattened by applying pressure from lead weights (6.75 kg) for 4 hours and dried under the fume hood for 1 hour. The tissue was dehydrated using four increasing concentrations of ethanol (75%, 95%, 100%, and 100%), 4 minutes each, followed by 20 minutes of xylene to render the tissue transparent. DEPEX mounting media (Electron Microscopy Sciences #13514, Hatfield, PA) was applied on the tissue and covered with a cover slip. The tissue was air-dried in the fume hood overnight. The total time for single labeling was 13 days.

Negative controls were prepared without any antibody or with secondary antibody alone. Antibody specificity was tested by *Abcam* and *Cell Signaling Technology* using Western Blot and immunofluorescence (https://www.abcam.com/products/primary-antibodies/cgrp-antibody-4901-ab81887.html; https://www.cellsignal.com/products/primary-antibodies/cgrp-d5r8f-rabbit-mab/14959). Western Blot analysis shows a single band at the correct molecular weight of CGRP in TT cells but not in HeLa cells. Moreover, confocal immunofluorescent analysis of TT and HeLa cells confirms that CGRP were in TT cells but not in HeLa cells. The specificity of both CGRP-antibodies used in this study was additionally confirmed by virtually identical staining patterns in the GI tract (more specifically in the stomach) of previous reports using other antibodies (*Peninsula Laboratories*: Hibbert et al. 2022, Smalilo et al. 2020, Spencer et al. 2016, *Sigma-Aldrich*: Tarif et al. 2021). The staining was shown to be reduced or completely abolished by preadsorption with CGRP from Peninsula (Domeneghini et al. 2004, Lennerz et al. 2008). Thus, we believe the CGRP antibodies used in this study are specific.

#### Dual-labeling immunofluorescence IHC

Flat-mount preparations were dual labeled with CGRP and either tyrosine hydroxylase (TH, sympathetic marker; dorsal n=2, ventral n=2) or vesicular acetylcholine transporter (VAChT, marker of excitatory enteric neurons, vagal pre-enteric neurons, and possibly gastric interneurons; dorsal n=2, ventral n=2). The listing of antibodies used in this study is provided in **Table 1.** The samples were first processed with primary and secondary antibodies for CGRP using the protocol as above for single labeling, then washed 6x5 minutes in 0.1 M PBS and blocked for 5 days using the blocking solution mentioned above before adding primary antibodies for TH or VAChT. The next steps for the second set of antibodies followed the same protocol for single labeling. The total time for double labeling was 26 days.

### 2.4 Fluoro-Gold (FG) counterstaining

To counterstain neurons in the myenteric plexus, three additional animals were used. Fluoro Gold (FG: 0.3 ml of 3mg/ml per mouse; Fluorochrome, Denver, CO) was injected intraperitoneally (i.p.) to counterstain neurons in the stomach and animals were perfused 3–5 days after FG injections. The stomachs were removed and peeled as described above for CGRP IHC labeling.

### 2.5 Tracer injection to the stomach wall

To verify the origins of CGRP-IR axons in the stomach, mice (n=4) were anesthetized with isoflurane (2-3%). The depth of anesthesia was determined by the lack of the hind paw and tail pinch withdrawal reflex. Tracer Fast DiI (Invitrogen, Catalog# D3899, Waltham, MA) was injected into the stomach wall using a Nanofil syringe microinjection system (NANOFIL-100, World Precision Instrument, Sarasota, FL). Each region, i.e., fundus, corpus, and antrum-pylorus received 3-4 injections (2 µL/each, 9-12 injections per surface) to cover the whole ventral or dorsal walls. The injection areas were cleaned thoroughly with cotton swabs, the displaced stomach was returned to the animal’s abdominal cavity in its original position, followed by closure of the abdominal muscle using interrupted sutures and the skin using a single continuous suture. The animals were returned to their cages for 14-16 days, then they were perfused. The left and right dorsal root ganglia (DRG, T1-T12) and left and right vagal nodose-petrosal ganglia complex (VNG) were removed for IHC labeling of CGRP.

### 2.6 Tissue scanning and montage assembly

The tissues were scanned with a Leica TCS SP5 Confocal Laser Scanning Microscope (Wetzlar, Germany). An argon-krypton (ArKr) laser (excitation wavelength at 488 nm and emission at 500-550 nm) was used to detect CGRP-IR axons and a helium-neon (HeNe) laser (excitation 543 nm) was used to detect TH-IR and VAChT-IR axons and background autofluorescence of the tissues (e.g., smooth muscles, blood vessels and myenteric ganglion cells). In addition, the 405 nm laser (excitation) was used to detect and verify FG-labeled myenteric ganglionic neurons. The scan was bidirectional and was achieved using a 20X oil immersion objective lens (N.A. 0.7, z-step of 1.5 μm), which yielded over 600 frames of confocal image stacks per montage. Each frame was 1024 x 1024 pixels. The image stacks were saved as .lif files and fully projected confocal images were saved as .tif files. All image tiles were stitched together with 30% overlap using Mosaic J and/or Adobe Photoshop to assemble the montage of the whole stomach (https://imagej.net/, RRID: SCR_003070; https://www.adobe.com/products/photoshop.html, RRID: SCR_014199). To better show the details of CGRP-IR axons and their connections to their gastric targets, regions of interest were scanned at a higher magnification using 40X (N.A. 1.25) and 100X (N.A. 1.4) oil immersion objective lens, 1.5X zoom, z-step of 1 μm. Using the confocal optical sections, we were able to detect what appear to be single axons. Modifications such as brightness and contrast adjustments, and scale bar additions were made using Adobe Photoshop.

The samples were also scanned with a Zeiss M2 Imager (Jena, Germany) that has an automatic stage and an Apotome 3 ZEN Digital Imaging system for Light Microscopy (https://www.zeiss.com/microscopy/en/products/software/zeiss-zen.html; RRID:SCR_013672) serves as the software program that controls the microscope settings. A preview image (5X lens, N.A. 0.16, excitation at 488 nm) was taken of the entire sample to define the borders. The preview image was then divided into multiple regions using the “Tile Region” tool. A higher magnification (20X lens, N.A. 0.8) was used to capture the z-stacks. Due to the uneven thickness across tissue, each tile had a different setting, and its z-stack range was set using the “Experimental Designer” tool. Following the scan, “raw” data was put through an automatic sequential batch process as follows: Apotome Deconvolution, Extended Depth of Focus (Max projection), Stitching, Image Export as .tiff. The exported regions were loaded in Photoshop to create a montage.

### 2.7 Morphological tracing and digitization

MBF Biosciences has developed the software Neurolucida 360® (https://www.mbfbioscience.com/neurolucida360, RRID:SCR_016788) to assist with the digitization, visualization, and quantification of neuronal structures. Another software by MBF Biosciences, Neurolucida Explorer® (https://www.mbfbioscience.com/neurolucida-explorer, RRID:SCR_017348), goes hand-in-hand with Neurolucida 360® to generate a full, comprehensive, quantitative analysis of reconstructed structures. We employed these tools to delineate the organization of the superficial vasculature and of CGRP-IR axons and terminals on both the dorsal and ventral surfaces of the stomach. Using the contour feature, we outlined the boundaries of well-preserved stomach whole mount tissues (fundus, corpus, antrum, pylorus, and intact esophagus), blood vessels (reconstructed in solid red) and individual axons (reconstructed in randomly assigned colors). Because digital reconstruction of these structures was performed on the stomach montages with over 300 maximum confocal projection images in the x-y plane that lacks z-plane information, the accuracy of tracing individual axons might be affected. We took into account the possibility of misidentifying and misinterpreting the directionality of the axons, therefore, a set of criteria was created to ensure the tracing method of axons was systematized (**Table 2**). By the same token, future investigations and studies that involve reconstruction of neuronal structures are made uniform using the same criteria.

### 2.8 Integration of axon tracing data into the 3D stomach scaffold

In order to render the reconstructed digital tracings in a more intuitive 3D configuration, we created a framework to transform the tracing data imaged from flat mount preparations to an anatomically accurate generic 3D mouse stomach scaffold. The generic mouse stomach scaffold was created using an open-source Python library called ScaffoldMaker (https://github.com/ABI-Software/scaffoldmaker, RRID:SCR_019003). This model was generated algorithmically from a set of anatomical and mathematical parameters derived from the standard shape and size of mouse stomachs. In line with the tissue preparation step, the whole stomach scaffold was split along its greater and lesser curvatures into its dorsal and ventral halves. Data mapping was described here using the ventral stomach scaffold, but this pipeline also applies for mapping to the dorsal stomach.

To bring the scaffold into the same coordinate space as the data, fiducial annotations were added to the stomach scaffold. The annotations used on the scaffold shared the same terminologies as those used for labeling the contours of the segmentation data. These fiducial annotations included the boundary of the distal end of the esophagus, fundus, limiting ridge, body, antrum, pylorus and the proximal end of the duodenum around the contours of the half stomach. Since consistent terms were used, annotations on the segmentation data could be matched to the same annotations on the stomach scaffold for alignment. Once the segmentation data was aligned with the stomach scaffold, the scaffold was then flattened and deformed so that the boundary of the scaffold matched the contour of the segmentation data. The location of the innervation and vasculature data was fitted into the flattened scaffold. Since the stomach scaffold was defined using a material coordinate system, the material coordinates of each data point derived from the segmentation data were calculated. These material coordinates effectively identified the position of any material (tissue) particle independent of its particular location in 3D space or how distorted the data was. This facilitated material embedding and provided a powerful reference frame to analyze and compare multiple sample data on one integrated scaffold. Using the mapped material locations, the flattened scaffold was then transformed together with its embedded innervation and vasculature data back to its 3D configuration.

The same protocol was applied to different samples, and this effectively consolidated data obtained from multiple samples in the same study onto one generic stomach scaffold, thereby providing a visualization aid for studying the spatial distribution and density of innervation and vasculature on the stomach.

## 3. Results

### 3.1 CGRP-IR axons and terminals in flat-mounts of the ventral and dorsal stomachs: an overview

The distribution and morphology of CGRP-IR axons and terminals in multiple layers at different regions of the flat-mount whole stomach were visualized in detail. Overall, CGRP-IR axons densely innervated the ventral and dorsal stomachs entirely. CGRP-IR varicose axons heavily wrapped around and ran in parallel with the blood vessels, formed varicose terminals in the myenteric ganglia, and innervated the circular and longitudinal muscles. Of note, blood vessels, muscles, and myenteric neurons were autofluorescent and they were present in the images scanned using 488 and 594 nm lasers as shown in Figure 4j-k, Figure 16d-e and Figure 16g-h. In addition, some unidentified cells also were autofluorescent and they were not innervated by CGRP-IR axons (Figure 10m).

**Figure 2** includes two montages of ventral and dorsal stomachs labeled with *Abcam’s* anti CGRP and **Figure 3** includes two montages of ventral and dorsal stomachs labeled with *Cell Signaling’s* anti-CGRP. Using our methodology, the *Cell Signaling* antibody had significantly less background noise than *Abcam’s*, therefore, the signal:noise ratio for the *Cell Signaling* antibody was much higher than that of the *Abcam* antibody. Nevertheless, when observed under higher magnification, both antibodies provided clear visualization of CGRP-IR axons and terminals around the blood vessels as well as in the muscles and myenteric ganglia. The detailed axons and varicosities are clearly visible in the stomach montage after zooming in.

**Figure 2.**
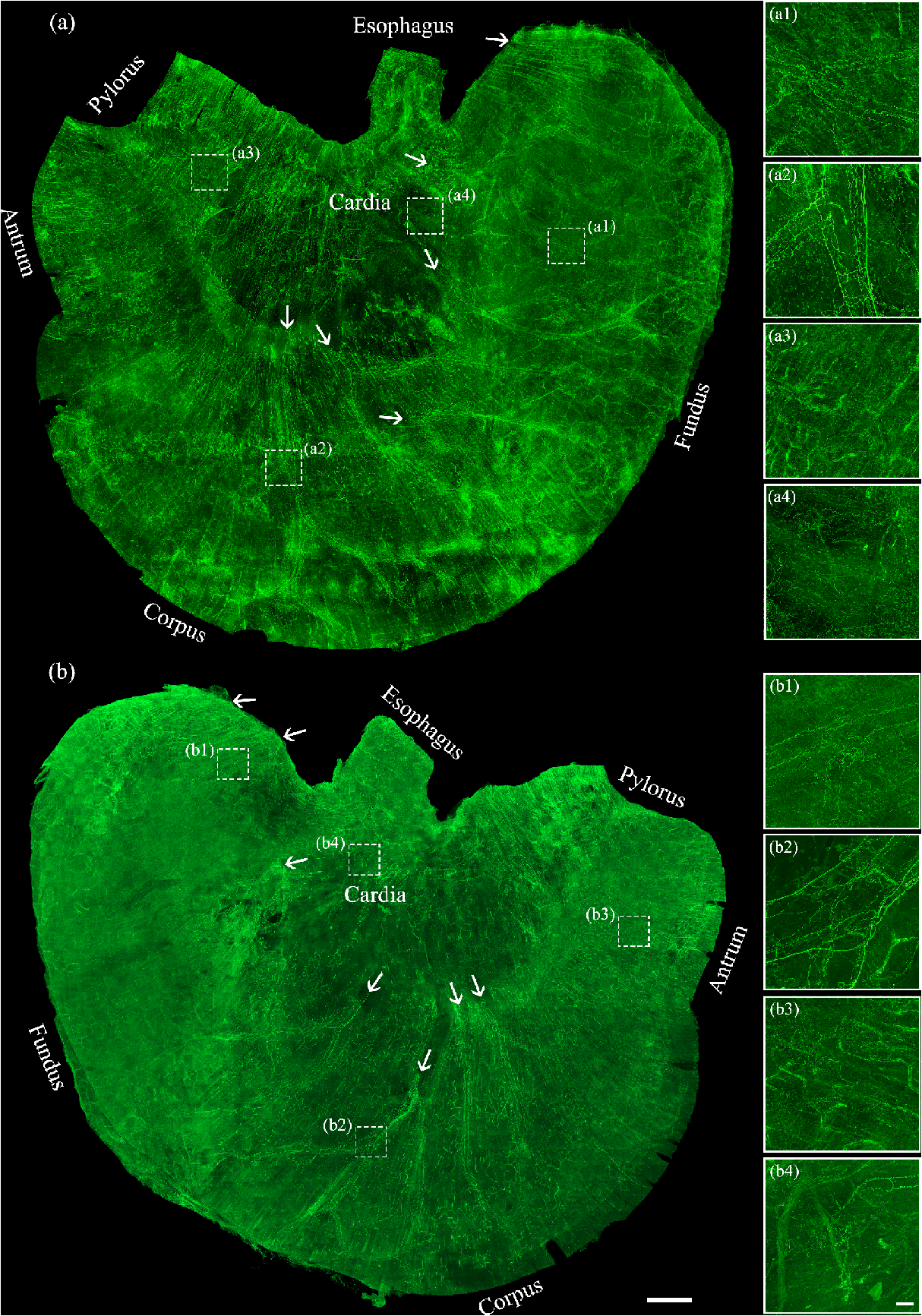
*Abcam* anti-CGRP: CGRP-IR axon innervation of stomach flat mounts. (a) Ventral. (b) Dorsal. Although the high background noise masked the CGRP signals in these montages, CGRP-IR axon distribution and terminal structures were clearly visible at higher magnification (a1-b4). The muscular layers of both ventral and dorsal stomach were innervated by CGRP-IR axons. The axon networks covered all regions including the fundus, corpus, antrum-pylorus, and cardia region. The cardia region was defined as the area immediately below the esophageal sphincter. Many CGRP-IR varicose axons traveled in the same directions as the longitudinal or circular muscles, ran in parallel with and wrapped around blood vessels, and formed varicose terminals around individual neurons in the myenteric ganglia (see the following figures for details). The arrows indicated the beginnings of some attached blood vessels. Scale bar = 1000 µm (montage), 100 µm (enlarged image). A higher resolution version of this figure is available in the Supporting Information.

**Figure 3.**
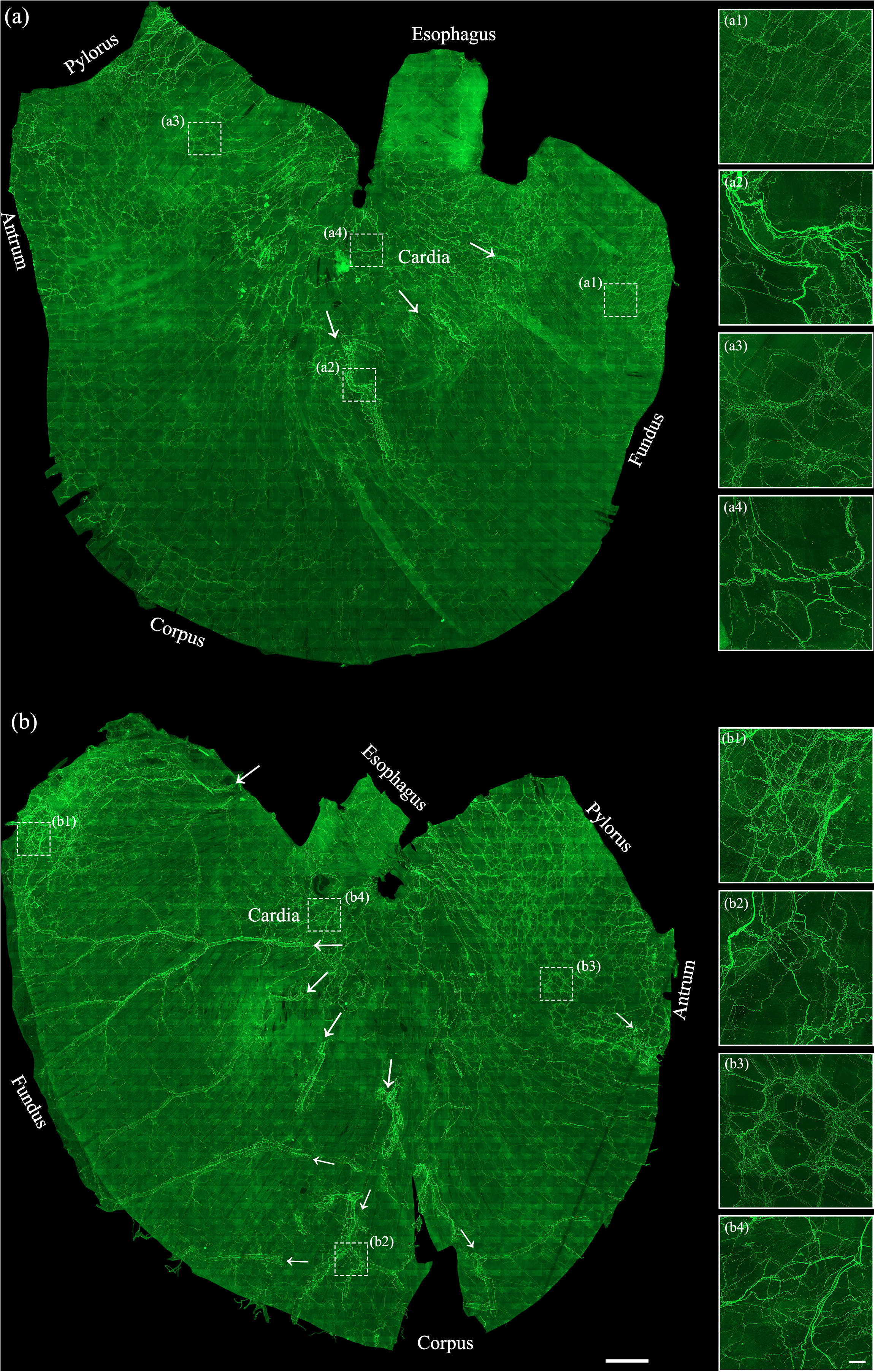
*Cell Signaling* anti-CGRP: CGRP-IR axon innervation of the stomach. (a) Ventral. (b) Dorsal. The background noise was much lower than that using Abcam’s anti-CGRP, and the innervation to the gastric targets was evident in these montages. The muscular layer of both ventral and dorsal surfaces of the stomach were innervated by CGRP-IR fibers. These axon networks covered all regions (a1-b4). Many CGRP-IR varicose axons traveled in the same direction as the circular and longitudinal muscles, ran in parallel with and wrapped around blood vessels, and formed varicose terminals around individual neurons in the myenteric ganglia. The arrows indicate the beginning of some attached blood vessels. Scale bar = 1000 µm (montage), 100 µm (enlarged image). A higher resolution version of this figure is available in the Supporting Information.

**Figure 4.**
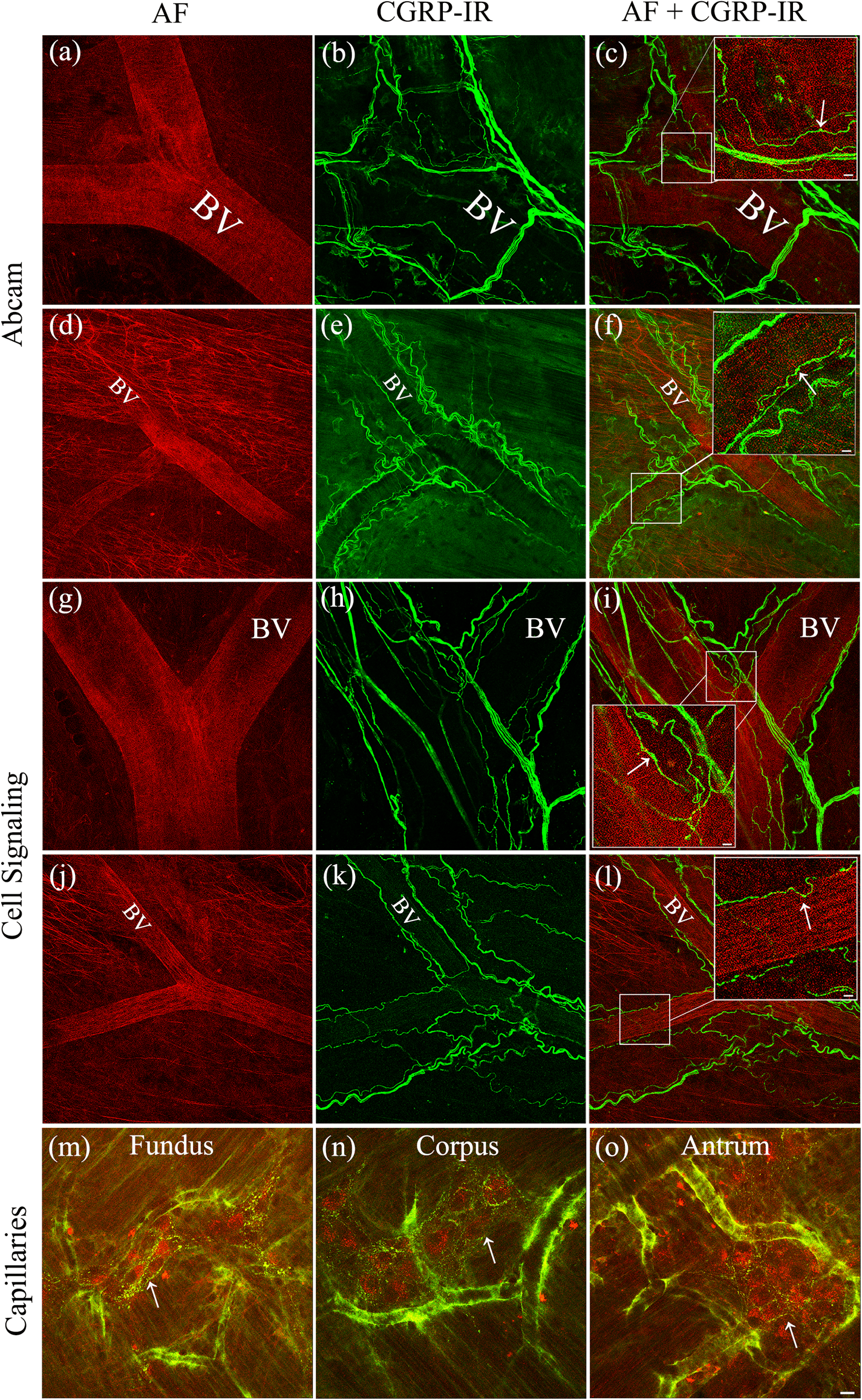
Single optical sections of CGRP-IR axon innervation along the blood vessels. (a, d, g, j) Autofluorescence of the arterioles (red) could be detected using the laser 543 nm. (b, e, h, k) CGRP-IR axon bundles bifurcated multiple times to either run in parallel with or wrap around the blood vessels. (c, f, i, l) While these axons were located at the immediate vicinity of the blood vessels, varicose axons with a smaller diameter (arrows inside insets) tended to be closer to the vessel wall. Compared to Abcam’s antibody (a-f), Cell Signaling produced similar quality of labeling around the blood vessels (g-l). (m-o) Single optical confocal images focused on the capillary bed in the fundus, corpus, and antrum of the stomach, respectively, of which CGRP-IR axons and terminals did not innervate. A good contrast of innervation would be the myenteric ganglia (arrows) near the capillary bed where CGRP-IR axons formed varicose contacts with individual neurons. AF = autofluorescence, BV = blood vessels. Scale bar in (o) = 20 µm and also applies to (a-n), in insets = 5 µm. A higher resolution version of this figure is available in the Supporting Information.

### 3.2 CGRP-IR axon innervation of the blood vessels

The most noticeable innervation from CGRP-IR axons was around the arteries. CGRP-IR axons entered the stomach alongside the major vasculature of the lesser curvature. Unlike the vagal afferent axons which entered the stomach from the lesser curvature with large nerve bundles (Fox et al. 2000, Wang and Powley 2000), we did not observe any large CGRP-IR axon bundles. The left and right gastric arteries extended from the lesser curvature to the greater curvature to cover most of the corpus of the ventral and dorsal stomach, respectively. In the antrum, we observed a small number of blood vessels coming from the right gastroepiploic artery. The fundus was also covered by small left gastroepiploic arterial branches. CGRP-IR fibers were abundant along the blood vessel walls and were present in all branches of the vasculature mentioned above. Innervation patterns revealed that the CGRP-IR axons were solid and bigger in diameter at the major branch of the blood vessels and became smaller after several bifurcations (and possibly trifurcations). These axons ran parallel with both sides of the blood vessel walls and often crossed to the other side of the wall to wrap around the vasculature (*Abcam*: **Figure 4a-f**, *Cell Signaling*: **Figure 4g-l**). Most fibers terminated when the blood vessels became capillaries (**Figure 4m-o**, shown by using autofluorescence). Some fibers terminated at other gastric targets such as the myenteric ganglia or the muscles.

### 3.3 CGRP-IR axons in the myenteric plexus

Shown in **Figure 5** is *Abcam*’s anti-CGRP and **Figure 6** is *Cell Signaling*’s anti-CGRP. CGRP IR axons and terminals were both present in the myenteric ganglia and the inter-ganglionic connectives of the ventral and dorsal stomach. It is evident that CGRP-IR axons formed exceptionally numerous varicose terminals around each individual myenteric neuron. These synaptic-like varicosities were more abundant in the corpus and antrum than in the fundus. While the myenteric neurons were surrounded by CGRP-IR nerve fibers, the cell bodies were not CGRP-IR, which led us to assume that the origin of these fibers was extrinsic and not intrinsic. In general, the CGRP-IR axon distribution and innervation pattern in the ventral and dorsal stomachs were similar. Furthermore, the antibodies from the two vendors (*Abcam* and *Cell Signaling*) yielded a similar pattern of innervation in the myenteric plexus.

**Figure 5.**
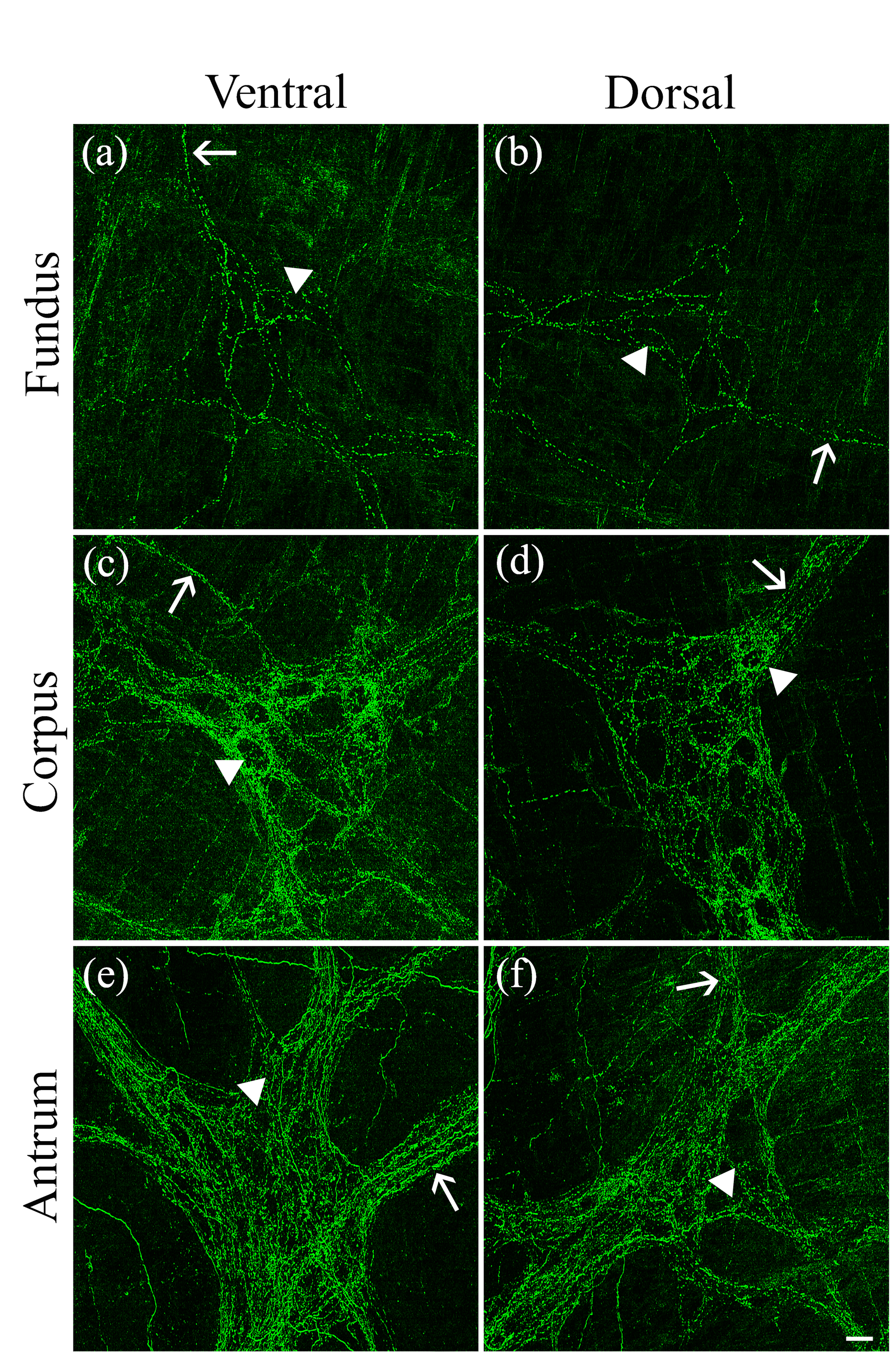
*Abcam* anti-CGRP: Partial projection confocal images revealed CGRP-IR axon innervation in the myenteric plexus at different regions of the stomach. (a, c, e) Ventral. (b, d, f) Dorsal. CGRP-IR axons traveled in the interganglionic connectives (arrows) and formed dense varicose terminals near and around many individual myenteric neurons (arrowheads; see FG labeled neurons in Figure 6). As a note: the autofluorescent background of the muscles was relatively higher than that of Cell Signaling. Scale bar = 20 µm. A higher resolution version of this figure is available in the Supporting Information.

**Figure 6.**
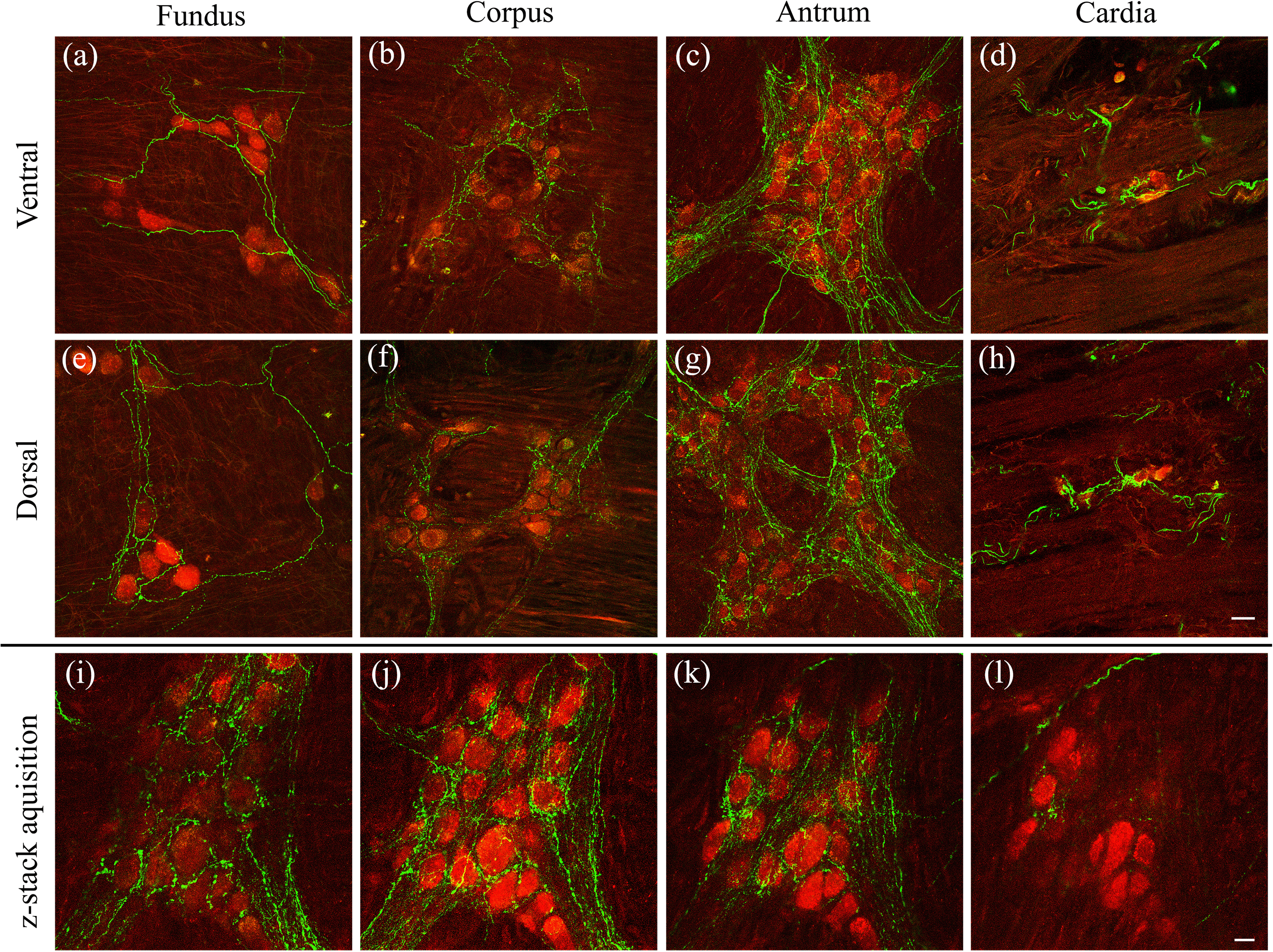
*Cell Signaling* anti-CGRP: Single optical sections revealed the myenteric ganglionic neurons stained with Fluoro-Gold (red). (a-d) Ventral. (e-h) Dorsal. CGRP-IR varicose axons (green) were present in the myenteric ganglia and the interganglionic connectives. (i-l) CGRP-IR varicose axons innervated individual myenteric neurons in the antrum region, presented in multiple confocal optical sections (z-step 1 μm). Scale bar in (h) = 20 µm and also applies to (a-g), in (l) = 10 µm and also applies to (j-l). A higher resolution version of this figure is available in the Supporting Information.

### 3.4 CGRP-IR axons and terminals in the longitudinal and circular muscles

**Figure 7** shows the CGRP-IR innervation of the muscular layer of the ventral and dorsal stomachs using *Cell Signaling* anti-CGRP. CGRP-IR fibers penetrated the muscular layer of both ventral and dorsal stomachs and ran in the directions of the longitudinal and circular muscles. They formed varicosities that were embedded in the muscle sheet and made varicose contacts with the muscles. The fibers in the muscles were found in all investigated regions including fundus, corpus, and antrum. These CGRP-IR fibers are also present abundantly in the cardia (*Abcam:* **Figure 8a,b**; *Cell Signaling:* **Figure 8c,d**).

**Figure 7.**
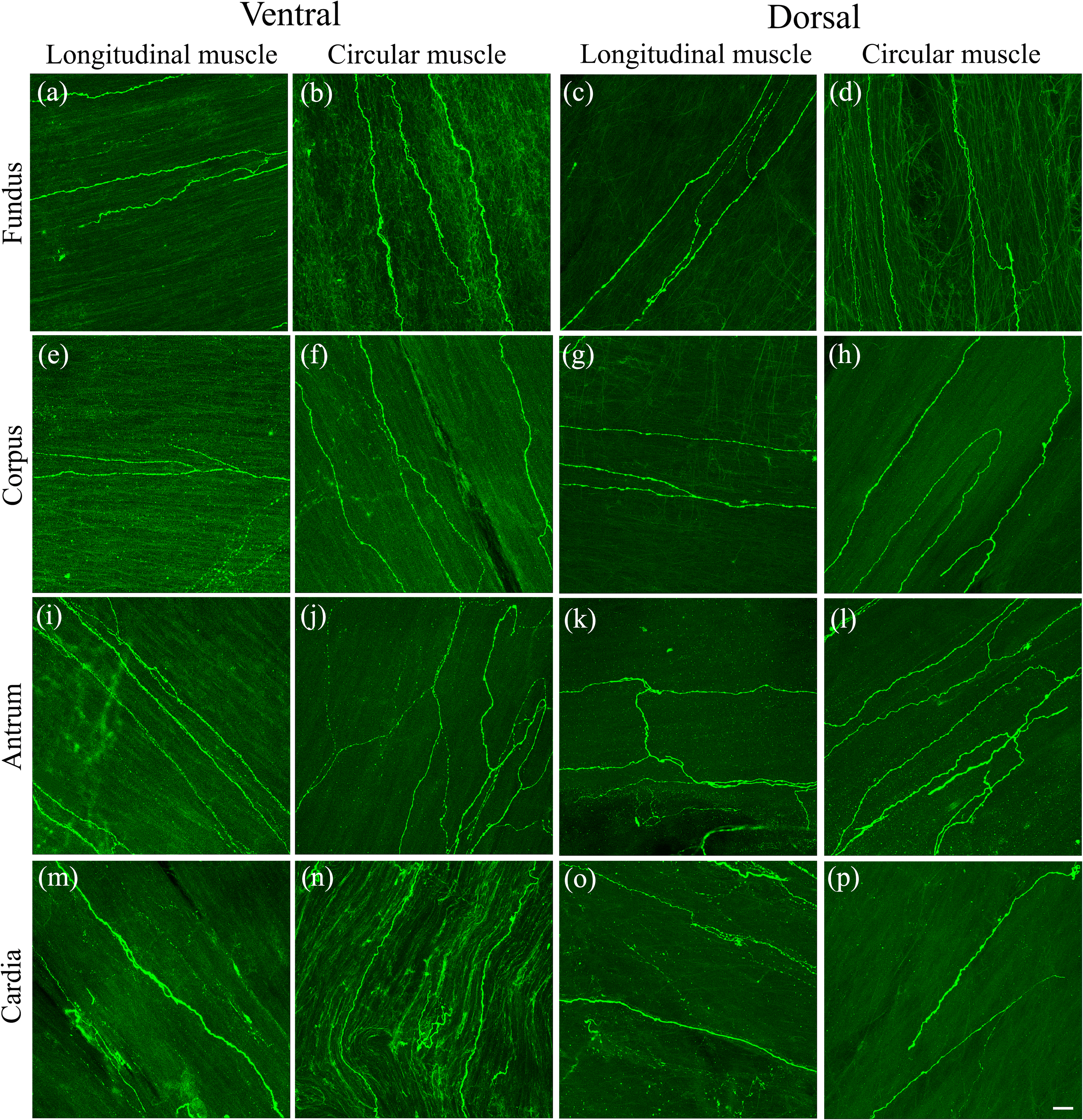
*Cell Signaling* anti-CGRP: Partial projection confocal images revealed CGRP-IR axon innervation at different locations of the stomach. (a, b, e, f, i, g, m, n) Ventral. (c, d, g, h, k, l, o, p) Dorsal. CGRP-IR fibers projected to both longitudinal and circular muscles and ran approximately parallel to the muscle fibers. Scale bar = 20 µm. A higher resolution version of this figure is available in the Supporting Information.

**Figure 8.**
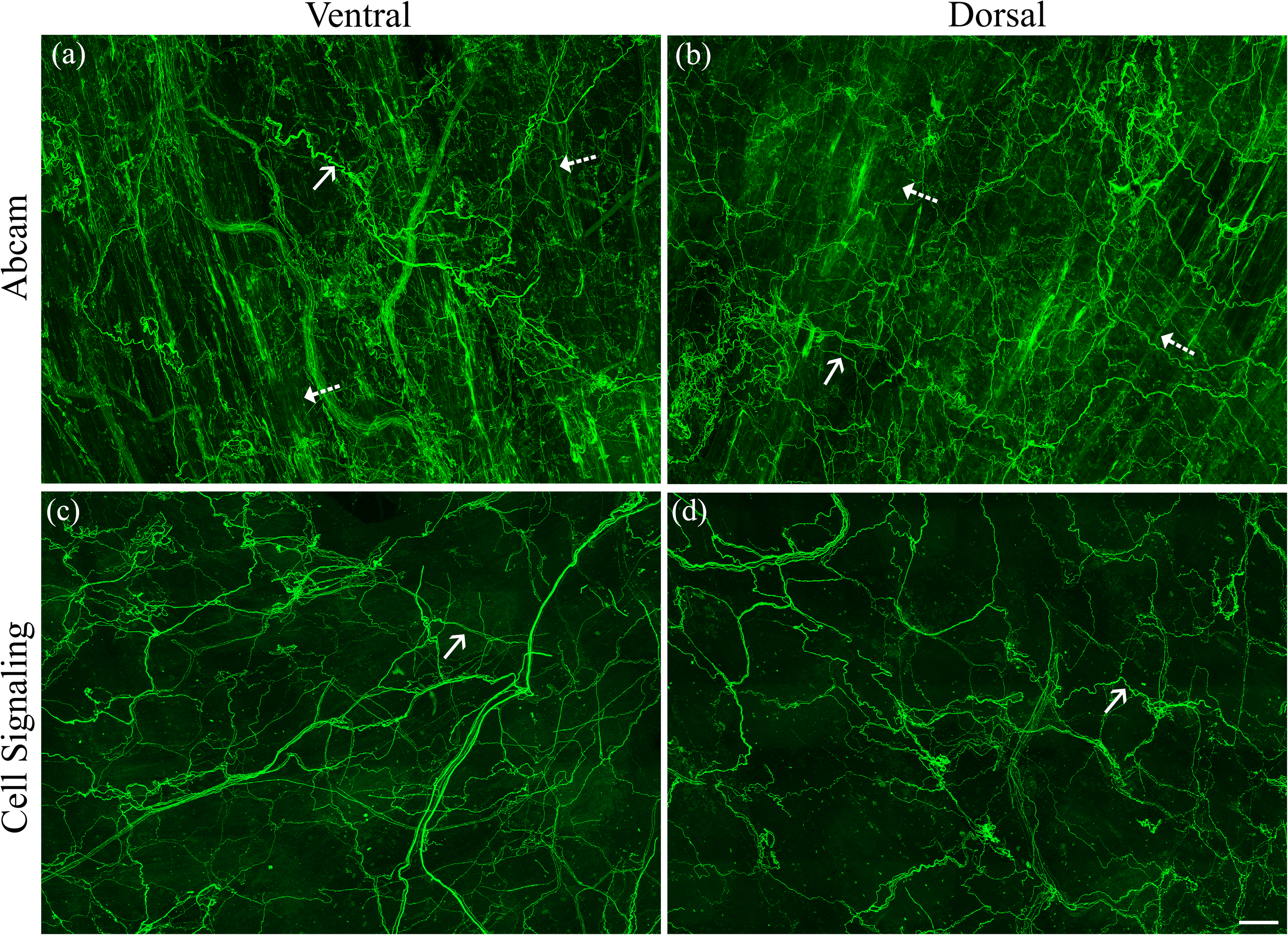
Maximum projection confocal images showed CGRP-IR axon innervation in the cardia region of the stomach. (a, b) Abcam. (c, d) Cell Signaling. These axons formed very complicated networks. Solid arrows point at CGRP-IR axons, dotted arrows point at autofluorescent background that might look like axons (mostly seen in Abcam stomachs). Scale bar = 100 µm. A higher resolution version of this figure is available in the Supporting Information.

**Figures 9 and 10** are a further investigation of CGRP-IR axon innervation of multiple layers of representative ventral and dorsal stomachs using *Cell Signaling* anti-CGRP. These axons first innervated the longitudinal muscle, traveled within the myenteric plexus, then innervated the circular muscle. While most axons in the muscular layers were parallel to the long axis of the longitudinal and circular muscles, some other axons, or parts of axons, did not follow the direction of the muscle fibers (autofluorescence highlights muscle direction), but rather crossed at an angle to the direction of the muscle cells. In general, the distribution pattern of CGRP-IR axons in the muscle walls of dorsal and ventral halves were similar.

**Figure 9.**
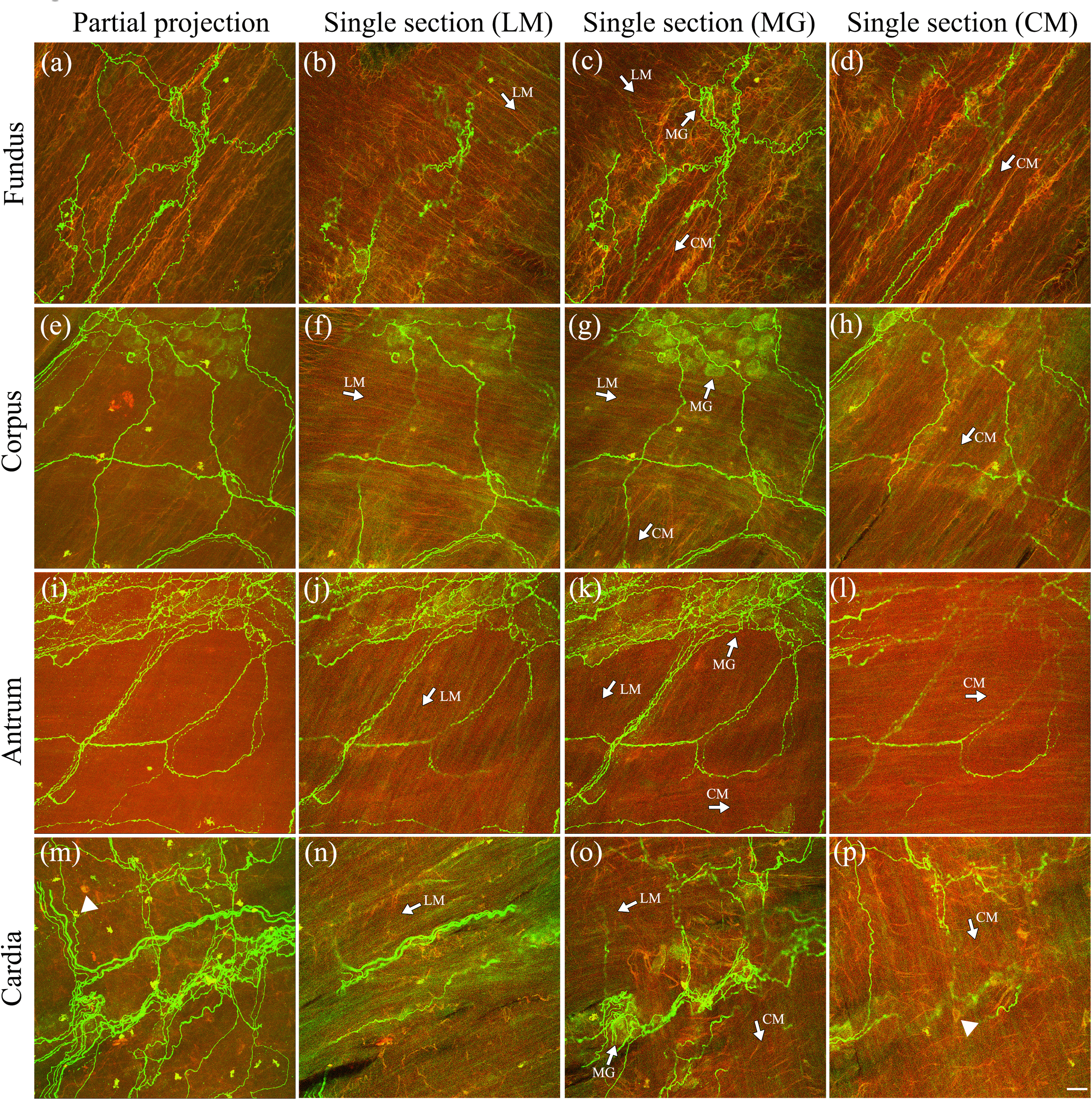
*Cell Signaling* anti-CGRP: Ventral stomach. Confocal images revealed the innervation from CGRP-IR varicose axons across multiple layers at different locations of the stomach. (a, e, i, m) Partial projections of the fundus, corpus, antrum and cardia, respectively. (b-d) Three single optical sections of (a) at different layers. (f-h) Three separated optical sections of (e). (j-l) Three separated optical sections of (i). (n-p) Three separated optical sections of (m). The CGRP-IR axons ran in parallel with the muscle fibers in the longitudinal layer (b, f, j, n), middle layer (myenteric plexus, c, g, k, o) and circular layer (d, h, l, p). White arrows indicate the direction of the muscles. White arrowheads point at autofluorescent cells. LM: longitudinal muscle, CM: circular muscle, MG: myenteric ganglia. Scale bar = 20 µm. A higher resolution version of this figure is available in the Supporting Information.

**Figure 10.**
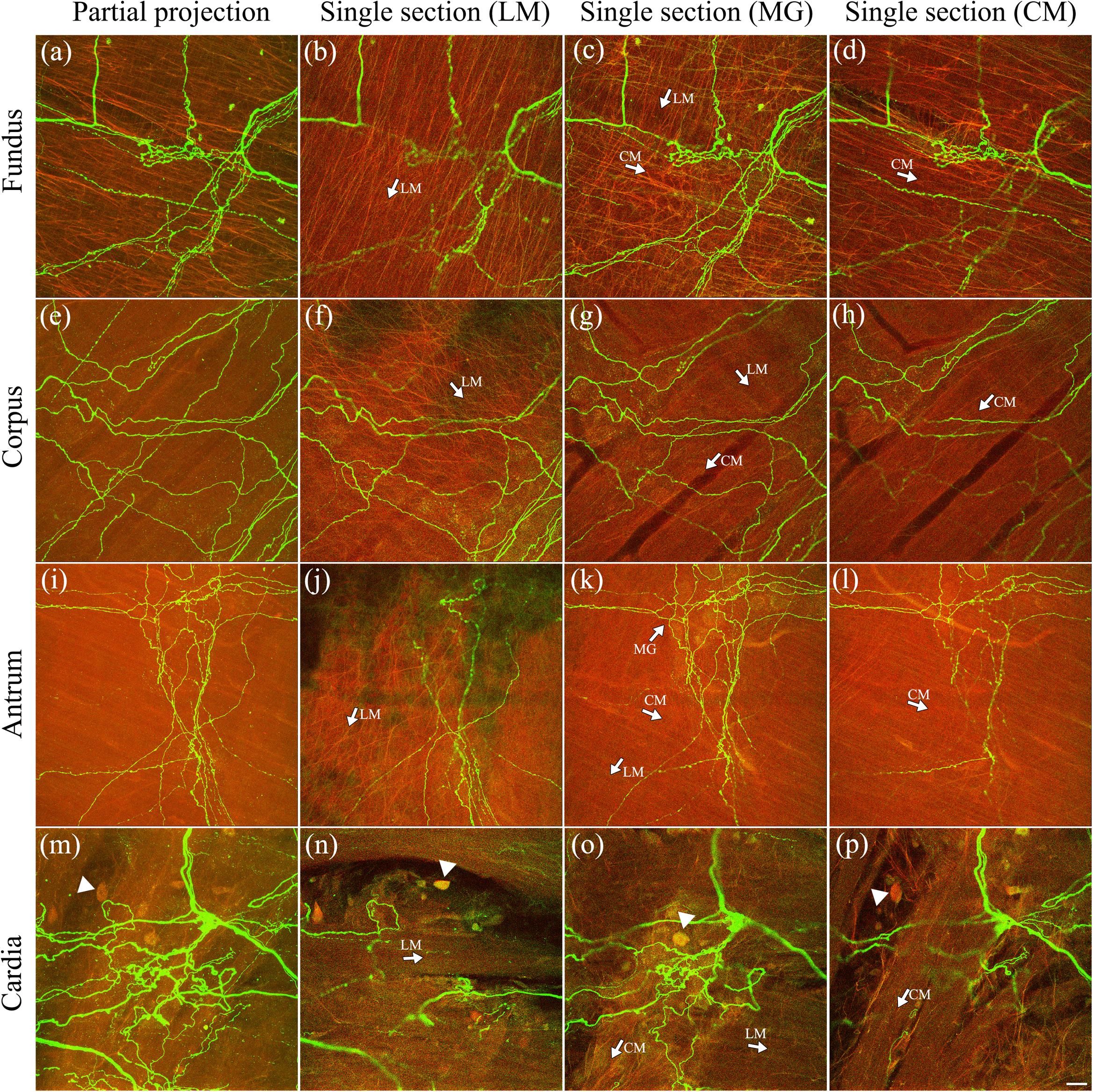
*Cell Signaling* anti-CGRP: Dorsal stomach. Single optical sections revealed the innervation from CGRP-IR varicose axons across multiple layers at different locations of the stomach. (a, e, i, m) Partial projections of the fundus, corpus, antrum and cardia, respectively(b-d) Three single optical sections of (a) at different layers. (f-h) Three separated optical sections of (e). (j-l) Three separated optical sections of (i). (n-p) Three separated optical sections of (m). The CGRP-IR axons ran in parallel with the muscle fibers in the longitudinal layer (b, f, j, n), middle layer (myenteric plexus, c, g, k, o) and circular layer (d, h, l, p). White arrows indicate the direction of the muscles. White arrowheads point at autofluorescent cells. LM: longitudinal muscle, CM: circular muscle, MG: myenteric ganglia. Scale bar = 20 µm. A higher resolution version of this figure is available in the Supporting Information.

### 3.5 Segmentation and integration of segmented data into a 3D stomach scaffold

To highlight the CGRP-IR axon innervation of the stomach, all CGRP-IR axons were traced using MBF Biosciences’ Neurolucida 360® software (**Figure 11**). The pattern of vasculature was similar in the ventral and dorsal stomachs (Ventral: **Figure 12a**, Dorsal: **Figure 13a**). The CGRP-IR axon innervation along the blood vessels, in the longitudinal and circular muscles, and in the myenteric ganglia were traced (Ventral: **Figure 12b-d**, Dorsal: **Figure 12b-d**). The nerve tracing of multiple animals resulted in a regional distribution pattern where a number of robust details were emphasized: 1) The CGRP-IR innervation was the most prominent in the vasculature and was mainly found in the corpus and fundus areas; 2) There were more CGRP-IR axons in the cardia and antrum-pylorus muscle layers than the fundus or corpus; 3) Axonal patterning was similar in dorsal and ventral stomachs.

**Figure 11.**
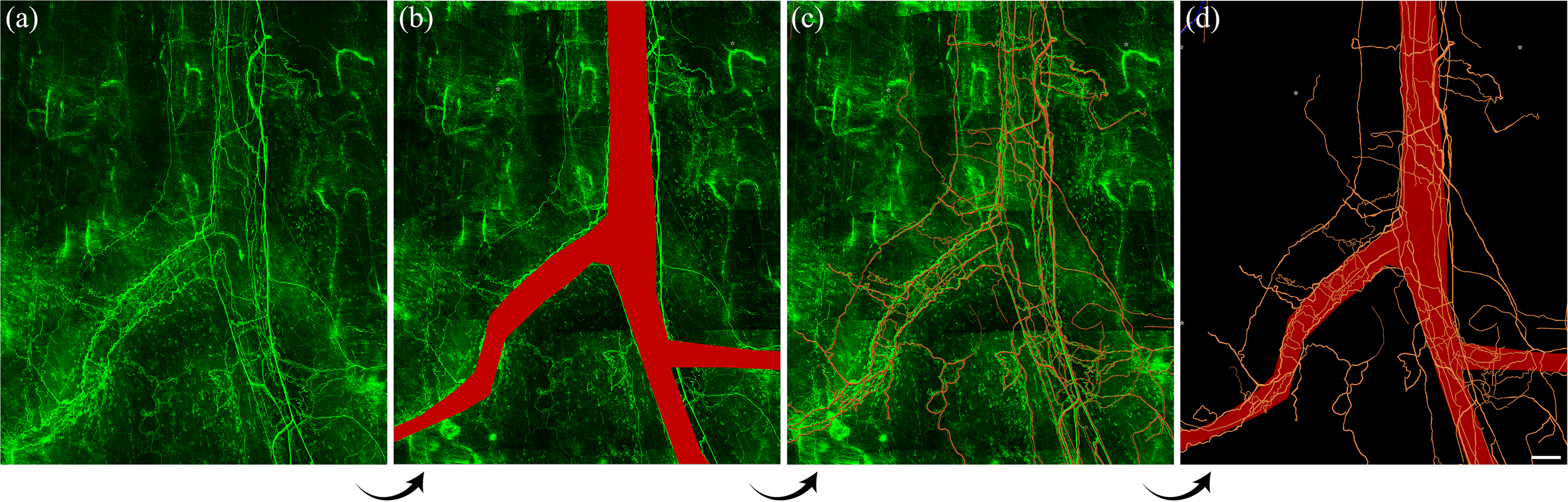
Workflow for tracing the vasculature and CGRP-IR axon innervation of the vasculature in the stomach using Neurolucida 360®. (a) is a representative blood vessel that was innervated by CGRP-IR axons. (b) A closed contour was made around the perimeter of the blood vessel using Contour mode. The contour was converted to a flat surface to display the area of the stomach of which the blood vessel occupied. Here, only the arteries were contoured. (c) The blood vessel contour was temporarily hidden, the axons were traced using Neuron Tracing mode (please see Table 2 for a set of criteria of how to classify and trace the axons). (d) The contouring and tracing data were stacked to create a complete tracing image while the confocal image was removed. Scale bar = 100 µm. A higher resolution version of this figure is available in the Supporting Information.

**Figure 12.**
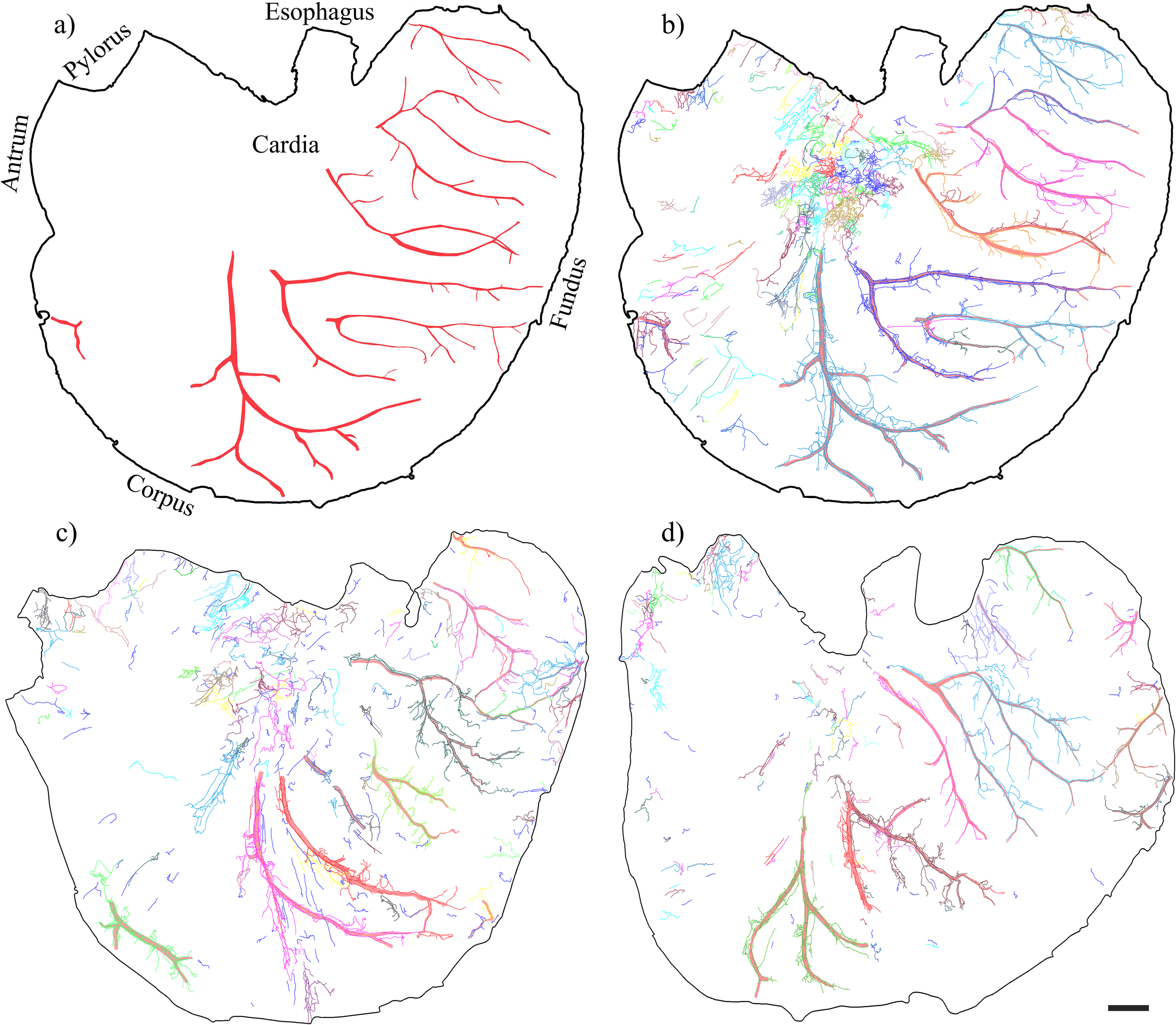
Reconstructed CGRP-IR axons of a ventral mouse stomach using Neurolucida 360. (a) Digitization of the vasculature displays the blood vessels’ distinctive distribution pattern. (b) CGRP-IR axons innervating the vasculature in stomach (a) as well as those in the muscular and myenteric plexus layers were identified and traced. These axons were assigned with a random color by the software. (c, d) Ventral stomachs of other animals also shared similar CGRP-IR axon innervation patterns. Scale bar = 1000 µm. A higher resolution version of this figure is available in the Supporting Information.

**Figure 13.**
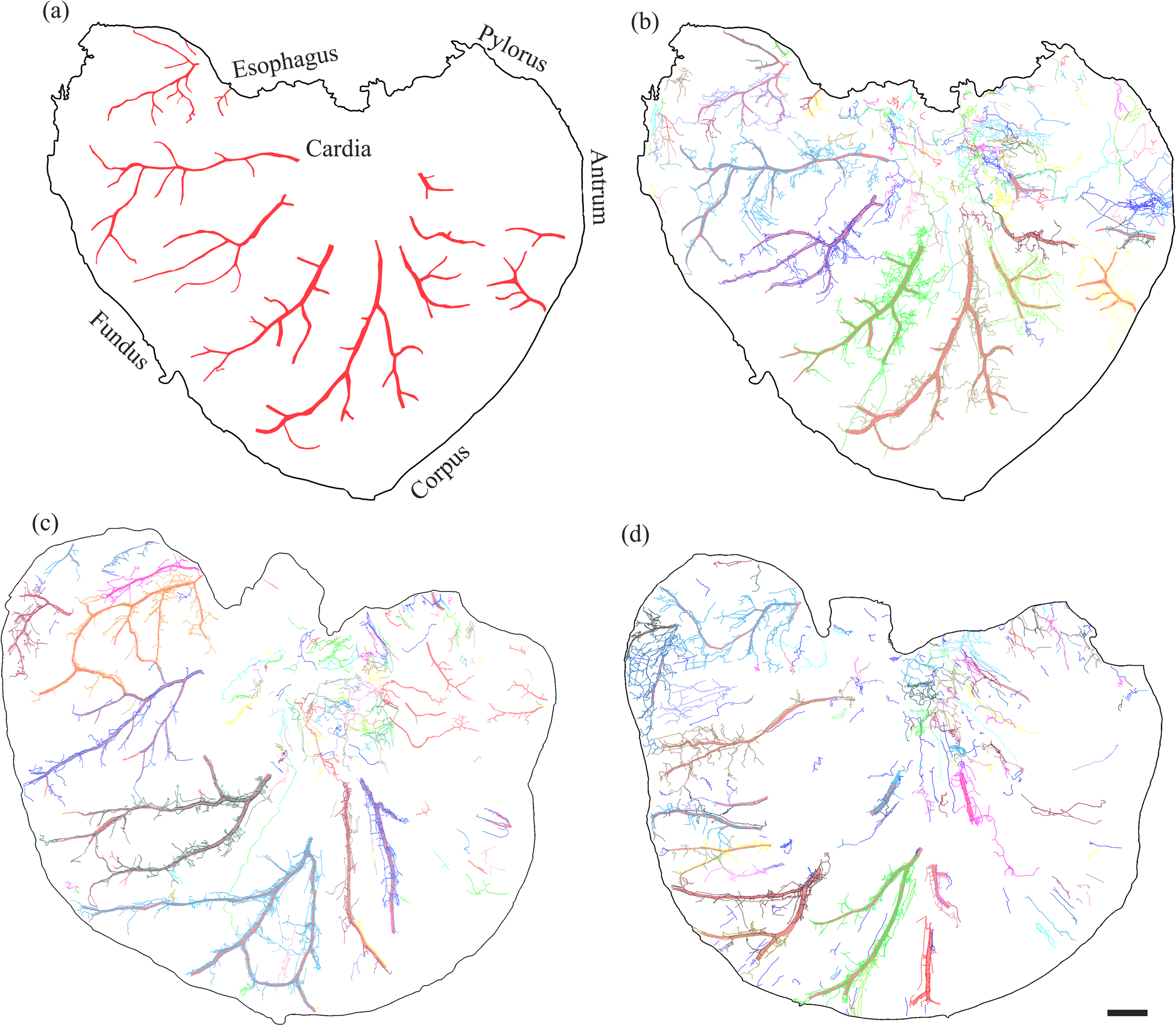
Reconstructed CGRP-IR axons of a dorsal mouse stomach using Neurolucida 360. (a) Digitization of the vasculature displays the blood vessels’ distinctive distribution pattern. (b) CGRP-IR axons innervating the vasculature in stomach (a) as well as those in the muscular and myenteric plexus layers were identified and traced. These axons were assigned with a random color by the software. (c, d) Dorsal stomachs of other animals also shared similar CGRP-IR axon innervation patterns. Scale bar = 1000 µm. A higher resolution version of this figure is available in the Supporting Information.

**Figure 14** highlights the end product of the innervation and vascular data being projected onto a mouse ventral stomach scaffold. The data, from being 2D, was brought into its 3D configuration as the flat scaffold is transformed back to the original 3D generic structure.

**Figure 14.**
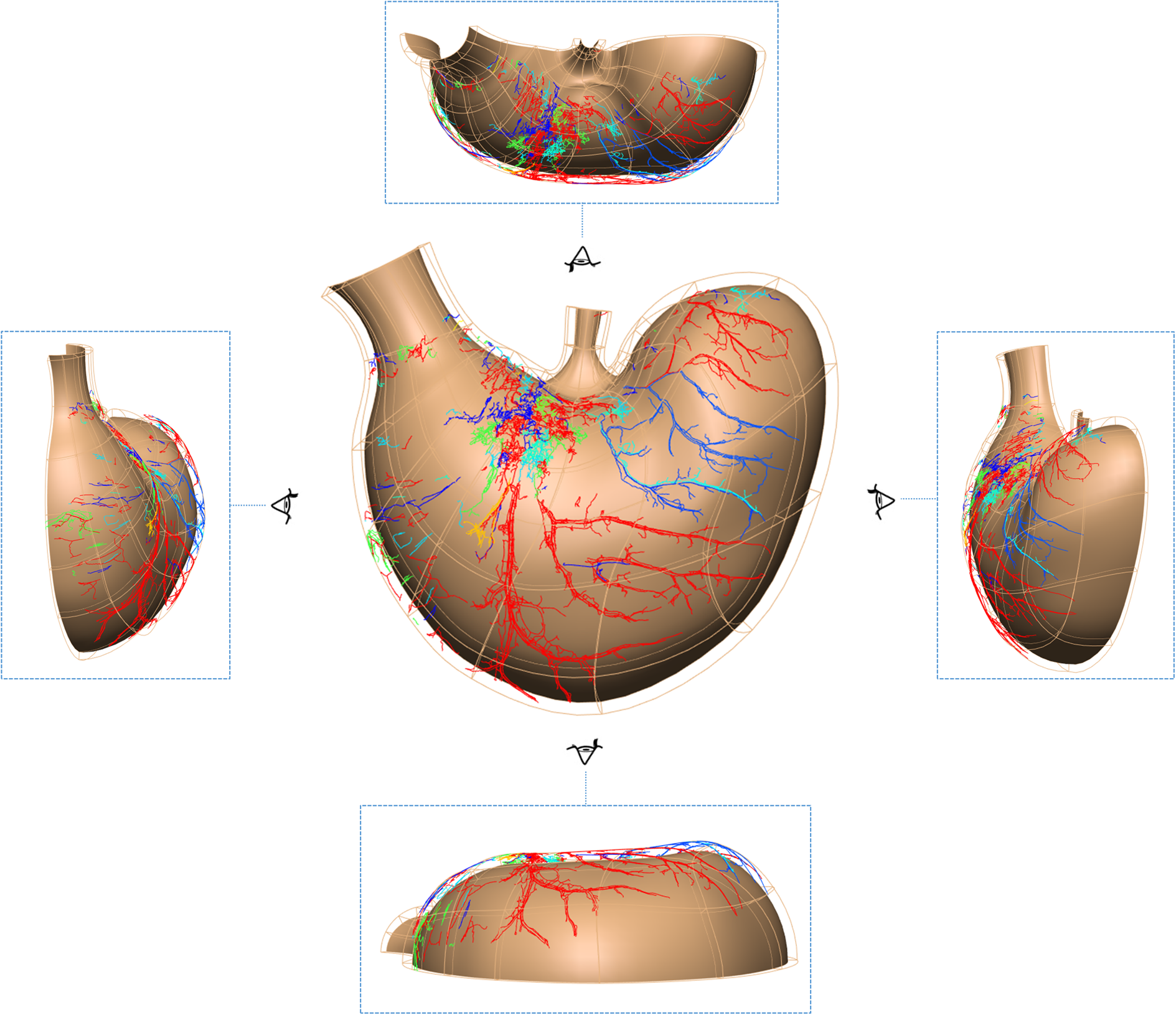
The integration of 2D innervation tracing and vascular contouring data into a 3D generic stomach scaffold. The data was mapped onto the material coordinate system of the flattened scaffold and embedded into the scaffold using fiducial markers to ensure alignment of the scaffold to the data, therefore, ensuring the accuracy of coordinate space. This allows the data to be brought into its 3D configuration as the flat scaffold is transformed back to the original 3D generic structure, as seen from different angles (front view - center, superior view - top, bilateral view - left+right, inferior view - bottom). A higher resolution version of this figure is available in the Supporting Information.

### 3.6 The origin of CGRP-IR axons in the stomach

As mentioned above, myenteric neurons were CGRP negative, suggesting that CGRP-IR axons were from extrinsic origin. To investigate the possible origins of CGRP-IR fibers in the stomach, we conducted three experiments:

1. After tracer DiI injection into the ventral and dorsal muscular walls, many neurons in the DRG and VNG were labeled. DiI-labeled neurons were dispersed in both the left and right T1 through T12 DRG and were most numerous in the T7 through T11 DRG (**Figure 15a-c**). IHC labeling with CGRP revealed that both the DRG and VNG contained double-labeled neurons with more double-labeled neurons in the DRG than VNG (**Figure 15d-f**), indicating that CGRP IR DRG afferent axons projected to the stomach more than the VNG did.

**Figure 15.**
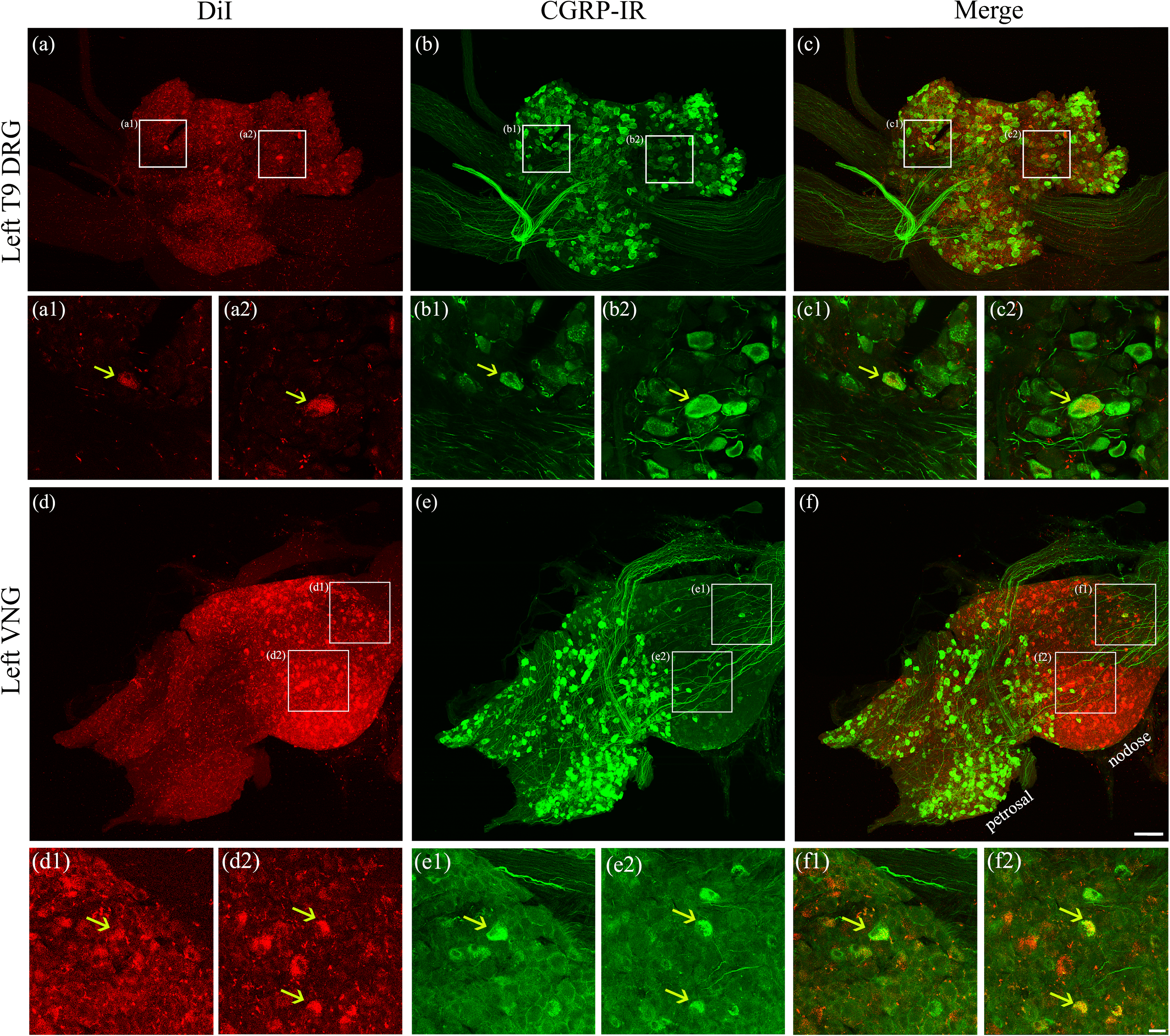
DiI injections into the stomach muscle wall retrogradely labeled the neurons within the left T9 DRG (a-c) and left vagal nodose + jugular ganglia complex (VNG) (d-f) followed by CGRP immunohistochemistry staining. (a-c) Maximum projection confocal images showed the double labeling of CGRP and DiI in multiple neurons within the ganglion. The number of dual labeled neurons peaked at T7 to T9 DRG. (d-f) Maximum projection confocal images showed the double labeling of CGRP and DiI of the neurons in the VNG, with the CGRP-IR neurons located mostly in the petrosal side of the VNG while most DiI-labeled neurons located in the nodose side of the VNG. (a1-f2) Single optical sections contained examples of double-labeled neurons at higher magnification, indicated by yellow arrows. Scale bar in (f) = 100 µm and also applies to (a-e), in (f2) = 20 µm and also applies to (a1-f1). A higher resolution version of this figure is available in the Supporting Information.

**Figure 16.**
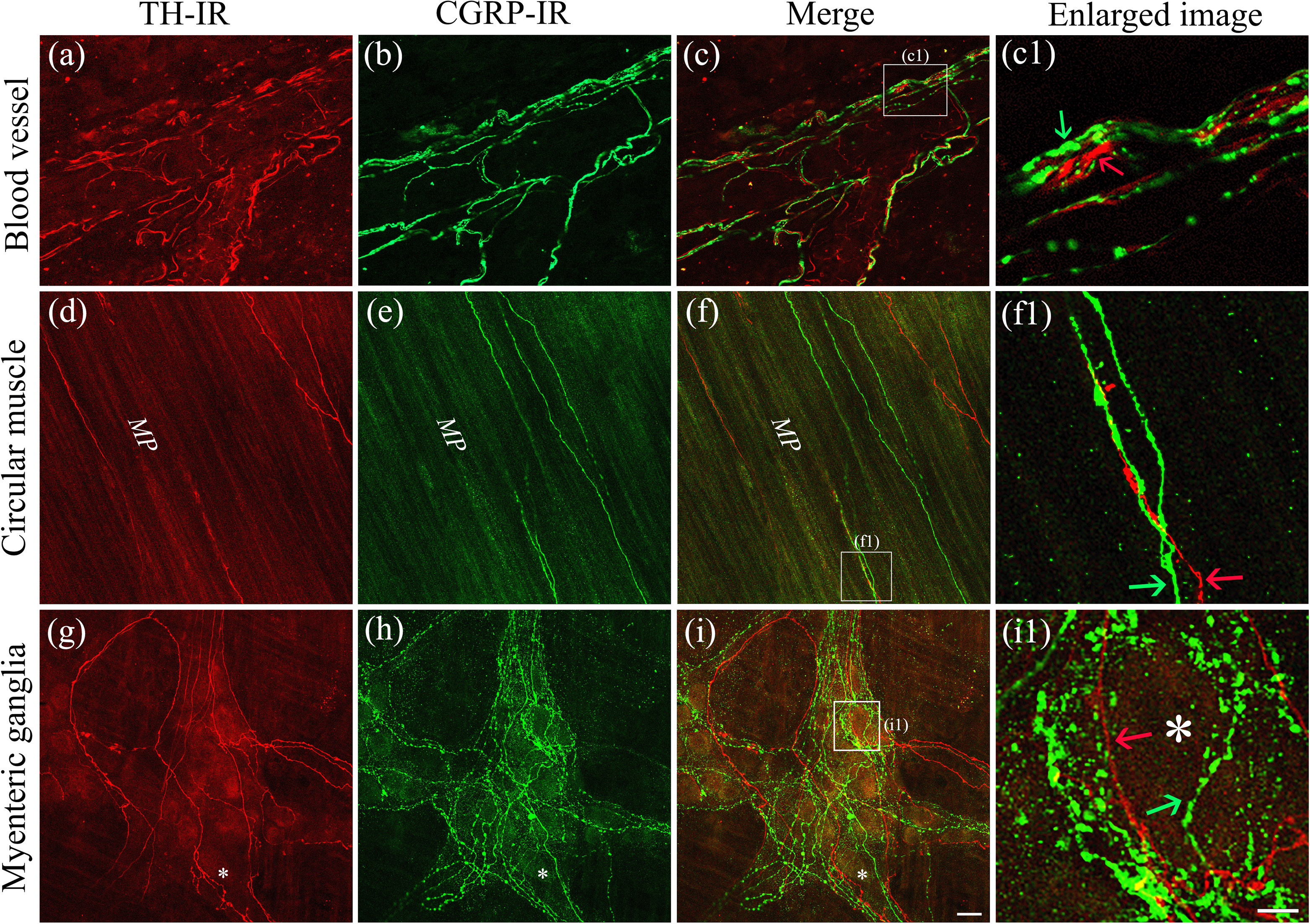
Dual labeling of TH (red) and CGRP (green) single optical sections showed the innervation of these two markers in different gastric targets. (a-c) A large number of non varicose TH-IR and varicose CGRP-IR axons ran parallel with each other along the blood vessels. (d-f) TH-IR and CGRP-IR axons projected in the direction of the circular muscle. (g-i) TH-IR axons innervated individual myenteric neurons by wrapping around the perimeter of the neurons while CGRP-IR axons deposited numerous varicosities around the neurons. (c1, f1, i1) High power magnification images of selected areas in (c, f, i), respectively, illustrated completely different innervation patterns of these two markers, indicating that colocalization did not exist. The red arrows point to fibers and terminals which are positive for TH but negative for CGRP, the green arrows point to axons and terminals which are positive for CGRP but negative for TH. Asterisks indicate the location of myenteric neurons. Scale bar in (i) = 20 µm and also applies to (a-h), (i1) = 5 µm and also applies to (c1, f1). A higher resolution version of this figure is available in the Supporting Information.

CGRP is a marker for sensory afferent axons from the spinal and vagal afferent neurons. However, whether CGRP could be of sympathetic/vagal efferent origin is not clear. Therefore, we double labeled this marker with TH and VAChT to examine whether CGRP might colocalize with TH and/or VAChT.

1. In **Figure 16**, the blood vessels, circular muscles, and myenteric ganglia of the stomach were both innervated by CGRP- and TH-IR fibers, which ran closely to each other, but no colocalization was detected. This indicates that CGRP-IR axons in the stomach were not of sympathetic origin.
2. Similarly, in **Figure 17**, VAChT-IR and CGRP-IR axons were both abundant along the blood vessels, in the circular muscles, and within the myenteric ganglia of the stomach. Their distribution patterns, however, were distinct, and no colocalization was detected. Since the two markers at the nerve terminals in the stomach did not colocalize, we concluded that CGRP-IR axons were not of vagal efferent origin.

**Figure 17.**
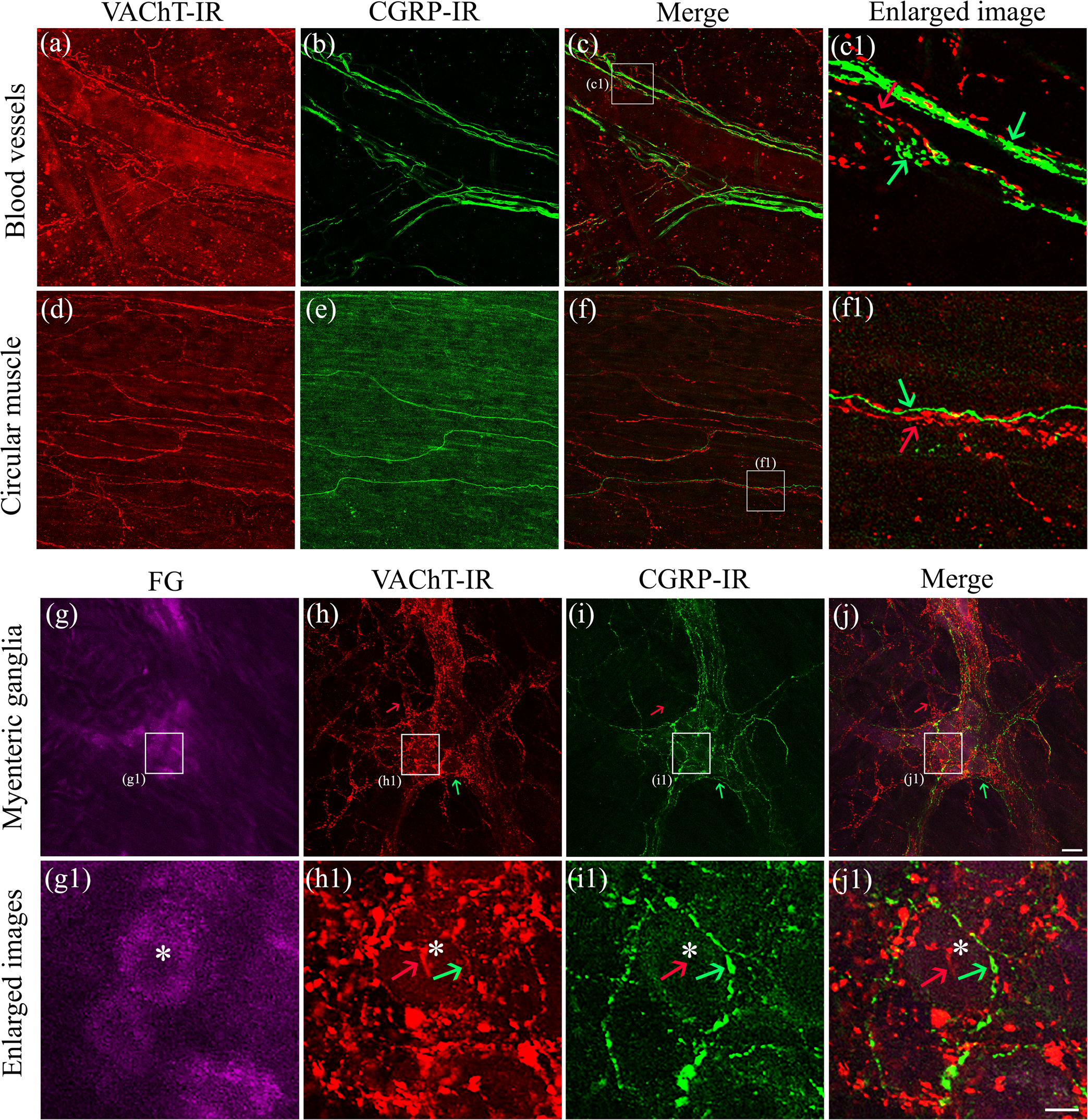
Dual labeling of VAChT (red) and CGRP (green) single optical sections showed the innervation of these two markers in different gastric targets. (a-c) A large number of varicose VAChT-IR and CGRP-IR axons ran parallel with each other along the blood vessels. (d-f) VAChT-IR axon innervation was much denser than that of CGRP-IR axons in the circular muscle. (g-j) VAChT-IR and CGRP-IR axons innervated individual myenteric neurons by wrapping around the neurons which were counterstained with Fluoro-Gold (FG, purple). (c1, f1, g1-j1) Enlarged images showed instances where VAChT-IR and CGRP-IR axons ran within close proximity, sometimes crossed over each other but were distinct in varicose thickness; hence, these fibers could be from the same nerve bundle but there was no colocalization. The red arrows point to fibers and terminals which are positive for VAChT but negative for CGRP, the green arrows point to axons and terminals which are positive for CGRP but negative for VAChT. Asterisks indicate the location of myenteric neurons. Scale bar in (j) = 20 µm and also applies to (a-i), in (j1) = 5 µm and also applies to (c1, f1, g1-i1). A higher resolution version of this figure is available in the Supporting Information.

## 4. Discussion

For the first time, we reported the organization and morphology of the CGRP-IR axons in flat mounts of the whole stomach muscular layers at the cellular/axonal/varicosity resolution. Strikingly, CGRP-IR axons ran along with and densely wrapped around the arteries. In addition, they innervated the myenteric neurons with varicose terminals which ran along/wrapped around individual neurons in the myenteric plexus at different regions of the whole stomach. In the longitudinal and circular muscles, CGRP-IR varicose axons were present, and they ran in the directions of the muscle fibers. Some axons crossed at an angle to the direction of the muscle cells. Overall, the CGRP-IR axons extensively innervated the whole stomach, and the innervation patterns were similar in both ventral and dorsal stomachs.

### 4.1 CGRP-IR axon innervation of the flat-mount whole stomach at the cellular/axonal/varicosity scale

We adapted several techniques, from tissue processing, immunohistochemical staining, image acquisition, image processing, to digitization of tracing data. Our study supports the work of others which showed CGRP-IR axon innervation of gastric targets and the origin of these axons, and it also has an important implication in delineating the comprehensive CGRP-IR gastric innervation while retaining the finest details. Through advanced imaging technologies, we were able to capture the details of innervation on specific targets and at different regions of the stomach. The novelty of this study lies in the creation of a comprehensive map where the continuity of the nerves can be preserved and the morphological progression of the axons at the targeted location such as the blood vessels, myenteric neurons, or the muscles is apparent. One of the most important highlights of the study is the visualization of the whole blood vessel trees and the abundance of CGRP-IR axons on the whole stomach’s arteries. We also observed a strong expression level of CGRP in the myenteric ganglia that implies that CGRP-IR axons may influence the myenteric ganglia to a greater degree than other gastric targets. Moreover, the CGRP-IR axon innervation pattern might be region specific, as certain areas such as the region immediately below the esophageal sphincter and the upper quadrant of fundus have denser innervation, and their biological significance still needs to be determined. Our findings provide a way to identify the density and pattern of innervation at specific areas and layers of the stomach. Thus, it can potentially offer some explanations to CGRP’s functions as well as provide vision for future physiological studies involving CGRP in the stomach.

Another novel technique worth mentioning here is the use of image digitization and automatic tracing of the signals. Regardless of all artifacts made by tears, cuts, or folds, the tracing enabled us to highlight the innervation, including the weak labeling of thin axons, and aided in visualizing valuable data. The digitization of tracing data also allowed us to select and filter the innervation based on the area/target of interest. On top of that, the 2D flat-mount CGRP-IR axon innervation tracing data, through computational modeling, was integrated into the 3D stomach scaffold. The integration of axon data into a generic scaffold gave us the ability to account for differences in the shapes and sizes of different tissue samples as well as compensate for the distortions due to tissue preparation, thereby allowing us to bring data from multiple samples into a single integrative scaffold for data comparison. The existence of such scaffolds enabled us to maintain the 3D nature of a multi-layered organ while accurately displaying each and every anatomical feature. Upon the completion of this study, not only was the CGRP-IR axon innervation data integrated into the scaffold, but other nociceptive nerve distribution data could also be prepared in a similar fashion. Having these types of information on a scaffold contributes to a more thorough understanding of the organ innervation, as well as making the data more accessible and replicable for other scientists. This is a joint effort from multiple researchers to not only produce the data, but also use the data to navigate 2D connectivity maps and build 3D organ scaffolds that can provide a comprehensive brain-organ atlas and “a common coordinate framework for cross-species comparisons” (Achanta et al. 2020, Leung et al. 2021, Osanlouy et al. 2021).

### 4.2 Origins of CGRP-IR axons

Historically, CGRP-IR neuron cell bodies existed in the myenteric plexus and submucosa of the GI tract such as the small intestine and colon of mouse (Furness et al. 2004, Mongardi Fantaguzzi et al. 2009, Qu et al. 2008, Tarif et al. 2021), colon of the rat (Mitsui 2009, Su et al. 1987), ileum of the lamb (Chiocchetti et al. 2005), and ileum of the guinea pig (Gibbins et al. 1985, Mawe et al. 1989). CGRP-IR neurons also existed in the stomachs of hamsters and pigs (Makowska 2019, Palus et al. 2018, Poomyachoti et al. 2002, Schochina et al. 1997). However, to our knowledge, we did not see any reports that CGRP-IR myenteric neurons were found in the mouse stomach. Consistent with literature (**Table 3**), our study could not detect positively labeled CGRP neurons in the stomach either, suggesting that the CGRP-IR axons might not come from intrinsic sources in the mouse stomachs or that the antibodies did not recognize CGRP precursors in the cell bodies.

Previous studies have shown that when extrinsic nerves projecting to the stomach were cut and time was allowed for the nerve endings to degenerate, all CGRP-IR nerve endings in the stomach were vastly reduced or disappeared completely. This effect was also observed after treatment with capsaicin (Furness et al 1991; 2001; Giancola et al. 2016, Green & Dockray 1988; Holzer 1998, Lee et al. 1987, Sternini et al. 1987, 1992, Suzuki et al. 1997). This indicates that CGRP IR fibers originated almost entirely from nerve cells located outside the stomach. Spencer and colleagues injected tracer Dextran-Biotin into the DRG to anterogradely label the spinal afferents in the large intestine, stomach, and bladder of mice (Spencer et al. 2014, 2016, 2018) and found that over 90% of the labeled spinal afferents were positive for CGRP, indicating that DRG is the major origin of the CGRP-IR axons in these visceral organs. Consistent with Spencer’s observations, we injected tracer DiI into the stomach wall to retrogradely label the neurons within the DRG and VNG found that a number of DiI-labeled neurons in the DRG across the thoracic vertebrae, especially T7 through T11, were also immunoreactive for CGRP. Compared to the total number of double-labeled neurons in the DRG, double-labeled VNG neurons seemed to be fewer. Along with previous studies, we conclude that the DRG were the major, and the VNG were the minor origin of CGRP-IR projections to the stomach (Holzer 1998, Rytel et al. 2015, Rytel and Calka 2016, Zhong et al. 2008).

Tracer injection into the VNG labeled the vagal afferents in the GI tract (Powley et al. 2016, 2019). These vagal afferents have terminal structures of intramuscular arrays (IMAs; stretch/length receptors), intraganglionic laminar ending (IGLEs; tension receptors) and mucosal innervating axons. IMAs appear in the muscle sheets as axons with a variety of varicosity depositions and IGLEs create flattened plates of contacts at the ganglionic surface in the mouse and rat stomachs (Fox et al. 2000, Castelluci et al. 2003, Powley et al. 2016, 2019). Our study found the potential structural similarity between the vagal afferents and CGRP-IR axons. However, the similarity is only limited to IMAs and CGRP-IR axons in the muscles. As for other gastric targets, CGRP-IR axons innervated the arteries of the whole stomach but vagal afferents did not. Also, CGRP-IR axons never formed IGLEs. These observations suggest that the CGRP DiI doubled labeled neurons in the VNG did project to the stomach, but most likely to the muscles only, and probably to the mucosa (e.g., Furness et al. 2001).

DmnX neurons send vagal preganglionic efferents to extensively innervate the myenteric ganglia of the rat stomach (Berthoud et al. 1990, Powley et al. 2019). CGRP expression was seen in the NTS, nucleus ambiguus (NA) neurons, and hypoglossal nucleus (XII) neurons but not in the DmnX neurons (Ladic and Buchan 1998, McGovern and Mazzone 2010). To further confirm CGRP-IR axons in the stomach do not come from DmnX, we used VAChT, a marker that labels vagal preganglionic efferents. The double labeling of VAChT and CGRP verified the presence of both VAChT-IR and CGRP-IR axons in the nerve terminals of the stomach, but no colocalization was detected. Therefore, we conclude that CGRP-IR axons in the stomach were not from vagal efferents.

The celiac and superior mesenteric ganglia project sympathetic postganglionic axons to the muscles and myenteric plexus of the stomach (Walter et al. 2016). In celiac ganglia, a dense population of CGRP-IR fibers passed by and occasionally surrounded the neurons (Kaestner et al. 2019). Nevertheless, no CGRP-positive cell bodies were found. Kaestner and colleagues in 2019 pointed out that the celiac neurons express a noradrenergic phenotype including tyrosine hydroxylase, neuropeptide Y, and vesicular monoamine transporters rather than sensory neuropeptides such as CGRP and SP. To further confirm that CGRP-IR axons in the stomach do not come from celiac ganglia, we used TH, a marker that specifically labels sympathetic postganglionic efferents. Double labeling of TH and CGRP showed that both TH-IR and CGRP IR axons were present in the stomach but did not colocalize on the blood vessels, muscles, and myenteric ganglia. Therefore, we conclude that CGRP-IR axons in the stomach were not sympathetic efferents.

To conclude, CGRP-IR axons in the stomach are of extrinsic origin, and the main supply of CGRP-IR axons in the stomach is the DRG, and to a lesser extent, the VNG.

### 4.3 Functional implications

CGRP synthesis in the peripheral sensory neurons and release from CGRP-IR nociceptors is achieved by the activation of TRPV1 (Clark and Gangula 2015, Luo et al. 2013, Meng et al. 2009). Upon delivery of harmful stimuli, afferent messages are sent through the dorsal horn of the spinal cord to the brain (Chen et al. 2010, Julius and Basbaum 2001). At the same time, the spinal cord receives sensory input and acts upon it by sending motor commands to the effector tissues. In addition, a local spread of signal, known as an axon reflex, occurs at the middle of the axonal processes and subserves a “dual sensory-efferent” function where afferent signals and neuropeptide release occur at the same nerve terminals (Holzer 2000, 2006, Szolcsanyi 2004, Yaprak 2008). The morphology and distribution of CGRP-IR nerve fibers suggest several biological activities that contribute to proper functioning of the stomach, including nociception, vasodilation, muscle relaxation, pro or anti-inflammatory responses (Bartho et al. 1991, 2008, Clark and Gangula 2015, Furness et al. 2014, Holzer 1991, 2006, Holzer and Farzi 2014, Kee et al. 2018, Kudrow et al. 2022, Timmermans et al. 1997). These nerve fibers also demonstrate a possible indirect role of CGRP in governing the gastric output, motility, secretion, as well as gastrointestinal hormone release in the stomach (Evangelista 2009, Kraenzlin et al. 1985).

### 4.4 CGRP in pathophysiological and pharmacological studies

Our bodies create a shift in neurochemical content of myenteric neural processes to accommodate for the damage caused by inflammation, gastric injury, or simply a change in environment (Schochina et al. 1997, Toole et al. 1999). Capsaicin-sensitive sensory nerves contain nociceptor markers such as CGRP and SP and are believed to be involved in local defense mechanisms against several conditions such as gastritis, erosions, ulcers, polyps, adenocarcinoma, and chronic inflammatory bowel diseases (Abdel-Salam et al. 1999, Dömötör 2005, Evangelista 2006, Ren et al. 2000).

A change in external environment factors (temperature, food, light) that causes food intake patterns to change can also affect the GI tract’s ability for storage, secretion, digestion, and absorption. For example, hibernation induced a significant increase in CGRP-IR cell bodies and nerve fibers projecting to the proximal stomach and proventriculus in hamsters (Schochina et al. 1997). This change is potentially a protective reflex mechanism to suppress active motility caused by substance P (Jensen 1997, Maton et al. 1988) and hypothermia-induced hypercontraction when food intake is limited and/or inactive periods are extended.

Despite the fact that CGRP can be useful as a potent vasodilator, muscle relaxant, or gastric secretion suppressor, we ought to acknowledge that too much production of this peptide can heighten our pain experience by increasing sensitivity and such overstimulation should not be prolonged (Julius and Basbaum 2001). Antagonists are, therefore, designed to reverse the nociceptive effects of CGRP on various tissues by downregulating the expression of CGRP in the DRG and sensory afferents (Bartho et al. 1991, Hou et al. 2020, Rudolf et al. 2005, Warzecha et al. 2001). Overall, the results from these studies can be used as a pivotal point to stimulate research into the development of new treatments for pathological conditions involving nociception.

## 5. Summary and future directions

We provided, for the first time, the organization and morphology of CGRP-IR axons in the flat mounts of the whole mouse stomach at the cellular/axonal/varicosity resolution. CGRP-IR axons were present in the longitudinal muscle, the myenteric plexus, and circular muscle. These axons were found in the fundus, corpus, antrum and cardia. The distribution and innervation pattern of the axons were similar in the dorsal and ventral stomachs.

This study offers a solid foundation for future quantitative analysis of CGRP-IR axon innervation in different targets at distinct locations of the stomach. The density analysis will provide a more definitive comparison in different regions of the gastric targets. The completion of this study has paved the way for potential experiments in the future where we can extensively investigate and compare the CGRP-IR axon innervation in the stomach between the two sexes. Moreover, our findings demonstrated that the DRG plays a major role in supplying CGRP-IR axons to the stomach, but whether the distribution and innervation pattern of CGRP-IR axons can completely represent the spinal afferent axons projecting to the stomach still needs to be further studied. Another important aspect to consider is the potential remodeling of CGRP-IR axons under pathological conditions and how it alters normal functions of the stomach. Since the generic stomach scaffold serves as a common coordinate framework, we are now able to integrate the CGRP-IR axon innervation data with different aspects of the stomach such as the musculature and vasculature data from other studies to investigate their interactions and create a comprehensive atlas containing anatomical, physiological, and molecular information. The integration of CGRP-IR axons and terminals into the generic stomach scaffold has laid out an anatomical foundation for future functional mapping of nociceptive axons, anatomical remodeling in pathological conditions and potential bioelectronic treatments for chronic pain.

## Acknowledgment

This study was supported by NIH 1 U01 HEAL/SPARC NS113867-01 and NIH 1R15HL137143-01A1. Many thanks to Kohlton Bendowski for editing the manuscript.

## Author’s contribution

Z.J.C. and T.L.P. designed the study. Ji.M. prepared and processed the tissues with immunohistochemistry. Ji.M. scanned samples using the confocal microscope, and Ja.M. scanned samples using Zeiss M2 Imager. D.N. performed tracing. M.H. and S.B. provided technical expertise on Neurolucida 360 usage. D.N. published the dataset on Pennsieve and S.B. curated the data. S.T. supervised M.H. and S.B. in MBF Bioscience software development. M.L. registered 3D mapping data onto the stomach scaffold and provided animated visualization of the scaffold. R.C. led the 3D scaffold developments. P.H. supervised M.L. and R.C. in MAP-CORE efforts for mapping data onto scaffolds. Ji.M., D.N., and A.B. assembled the figures. D.N and Ji.M. wrote the manuscript with supervision from Z.J.C., J.B.F. and T.L.P. J.C. helped the study overall. A.B. helped revise the manuscript. All authors contributed to the article and approved the submitted version.

## Data Availability Statement

The dataset used and analyzed in this study was uploaded and published (https://sparc.science/datasets/230?type=dataset) and will be available upon request from the corresponding author.

## References

1. Abdel-Salam OME, Debreceni A, Mózsik G, Szolcsányi J. (1999) Capsaicin-sensitive afferent sensory nerves in modulating gastric mucosal defense against noxious agents. Journal of Physiology-Paris, 93(5), 443–454. https://doi.org/10.1016/S0928-4257(99)00115-1.

2. Achanta S, Gorky J, Leung C, Moss A, Robbins S, Eisenman L, Chen J, Tappan S, Heal M, Farahani N, Huffman T, England S, Cheng ZJ, Vadigepalli R, Schwaber JS. (2020) A Comprehensive Integrated Anatomical and Molecular Atlas of Rat Intrinsic Cardiac Nervous System. iScience, 23(6):101140. doi: 10.1016/j.isci.2020.101140.

3. Bartho L, Benko R, Holzer-Petsche U, Holzer P, Undi S, Wolf M. (2008) Role of extrinsic afferent neurons in gastrointestinal motility. Eur Rev Med Pharmacol Sci, 12 Suppl 1:21–31.

4. Barthó L, Kóczán G, Holzer P, Maggi CA, Szolcsányi J. (1991) Antagonism of the effects of calcitonin gene-related peptide and of capsaicin on the guinea-pig isolated ileum by human alpha-calcitonin gene-related peptide(8-37). Neurosci Lett, 129(1):156–9. doi: 10.1016/0304-3940(91)90744-e.

5. Bell D, McDermott BJ. (1996) Calcitonin gene-related peptide in the cardiovascular system: characterization of receptor populations and their (patho)physiological significance. Pharmacol Rev, 48(2):253–88.

6. Berthoud HR, Carlson NR, Powley TL. (1991) Topography of efferent vagal innervation of the rat gastrointestinal tract. Am J Physiol, 260(1 Pt 2):R200–7. doi: 10.1152/ajpregu.1991.260.1.R200.

7. Brain SD, Grant AD. (2004) Vascular actions of calcitonin gene-related peptide and adrenomedullin. Physiol Rev, 84(3):903–34. doi:10.1152/physrev.00037.2003.

8. Castelucci P, Robbins HL, Furness JB. (2003) P2X(2) purine receptor immunoreactivity of intraganglionic laminar endings in the mouse gastrointestinal tract. Cell Tissue Res, 312(2):167–74. doi: 10.1007/s00441-003-0715-3.

9. Chen LJ, Zhang FG, Li J, Song HX, Zhou LB, Yao BC, Li F, Li WC. (2010) Expression of calcitonin gene-related peptide in anterior and posterior horns of the spinal cord after brachial plexus injury. J Clin Neurosci, 17(1):87–91. doi: 10.1016/j.jocn.2009.03.042.

10. Chiocchetti R, Grandis A, Bombardi C, Lucchi ML, Dal Lago DT, Bortolami R, Furness JB. (2006) Extrinsic and intrinsic sources of calcitonin gene-related peptide immunoreactivity in the lamb ileum: a morphometric and neurochemical investigation. Cell Tissue Res, 323(2):183–96. doi: 10.1007/s00441-005-0075-2.

11. Clark SY, Gangula PR. (2015) Role of Calcitonin Gene-related Peptide in Gastric Motility Function: Animal and Human Studies. J Gastrointest Dig Syst 5:276. doi:10.4172/2161-069X.1000276.

12. Di Natale MR, Patten L, Molero JC, Stebbing MJ, Hunne B, Wang X, Liu Z, Furness JB. (2021) Organization of the musculature of the rat stomach. J Anat. doi: 10.1111/joa.13587.

13. Domeneghini C, Radaelli G, Arrighi S, Bosi G, Dolera M. (2004) Cholinergic, nitrergic and peptidergic (Substance P- and CGRP-utilizing) innervation of the horse intestine. A histochemical and immunohistochemical study. Histol Histopathol, 19(2):357–70. doi: 10.14670/HH-19.357.

14. Dömötör A, Peidl Z, Vincze A, Hunyady B, Szolcsányi J, Kereskay L, Szekeres G, Mózsik G. (2005) Immunohistochemical distribution of vanilloid receptor, calcitonin-gene related peptide and substance P in gastrointestinal mucosa of patients with different gastrointestinal disorders. Inflammopharmacology,13(1-3):161–77. doi: 10.1163/156856005774423737.

15. Ekblad E, Ekelund M, Graffner H, Håkanson R, Sundler F. (1985) Peptide-containing nerve fibers in the stomach wall of rat and mouse. Gastroenterology, 89(1), 73–85. doi:10.1016/0016-5085(85)90747-4.

16. Evangelista S. (2006) Role of Sensory Neurons in Restitution and Healing of Gastric Ulcers, Current Pharmaceutical Design, 12(23). doi:10.2174/138161206777947632.

17. Evangelista S. (2009) Role of calcitonin gene-related Peptide in gastric mucosal defense and healing. Curr Pharm Des, 15(30):3571–6. doi:10.2174/138161209789207024.

18. Eysselein VE, Reinshagen M, Patel A, Davis W, Nast C, Sternini C. (1992) Calcitonin gene related peptide in inflammatory bowel disease and experimentally induced colitis. Ann N Y Acad Sci, 657:319–27. doi: 10.1111/j.1749-6632.1992.tb22779.x

19. Fox EA, Phillips RJ, Martinson FA, Baronowsky EA, Powley TL. (2000) Vagal afferent innervation of smooth muscle in the stomach and duodenum of the mouse: Morphology and topography. J Comp Neurol, 428: 558–576. https://doi.org/10.1002/1096-9861(20001218)428:3<558::AID-CNE11>3.0.CO;2-M

20. Frias B, Merighi A. (2016) Capsaicin, Nociception and Pain. Molecules, 21(6):797. doi: 10.3390/molecules21060797.

21. Furness JB, Lloyd KC, Sternini C, Walsh JH. (1991) Evidence that myenteric neurons of the gastric corpus project to both the mucosa and the external muscle: myectomy operations on the canine stomach. Cell Tissue Res, 266(3):475–81. doi: 10.1007/BF00318588.

22. Furness JB, Koopmans HS, Robbins HL, Clerc N, Tobin JM, Morris MJ. (2001) Effects of vagal and splanchnic section on food intake, weight, serum leptin and hypothalamic neuropeptide Y in rat. Auton Neurosci ;92(1-2):28–36. doi: 10.1016/S1566-0702(01)00311-3.

23. Furness JB, Robbins HL, Xiao J, Stebbing MJ, Nurgali K. (2004) Projections and chemistry of Dogiel type II neurons in the mouse colon. Cell Tissue Res, 317(1):1–12. doi: 10.1007/s00441-004-0895-5.

24. Furness JB, Callaghan BP, Rivera LR, Cho HJ. (2014) The enteric nervous system and gastrointestinal innervation: integrated local and central control. Adv Exp Med Biol, 817:39–71. doi: 10.1007/978-1-4939-0897-4_3.

25. Furness JB, Di Natale M, Hunne B, Oparija-Rogenmozere L, Ward SM, Sasse KC, Powley TL, Stebbing MJ, Jaffey D, Fothergill LJ. (2020) The identification of neuronal control pathways supplying effector tissues in the stomach. Cell Tissue Res, 382(3):433–445. doi: 10.1007/s00441-020-03294-7.

26. Giancola F, Gentilini F, Romagnoli N, Spadari A, Turba ME, Giunta M, Sadeghinezhad J, Sorteni C, Chiocchetti R. (2016) Extrinsic innervation of ileum and pelvic flexure of foals with ileocolonic aganglionosis. Cell Tissue Res, 366(1):13–22. doi: 10.1007/s00441-016-2422-x.

27. Gibbins IL, Furness JB, Costa M, MacIntyre I, Hillyard CJ, & Girgis S. (1985) Co-localization of calcitonin gene-related peptide-like immunoreactivity with substance P in cutaneous, vascular and visceral sensory neurons of guinea pigs. Neuroscience Letters, 57(2), 125–130. doi:10.1016/0304-3940(85)90050-3.

28. Godlewski J, Kaleczyc J. (2010) Somatostatin, substance P and calcitonin gene-related peptide positive intramural nerve structures of the human large intestine affected by carcinoma. Folia Histochem Cytobiol, 48(3):475–83. doi:10.2478/v10042-010-0079-y.

29. Green T, Dockray GJ. (1988) Characterization of the peptidergic afferent innervation of the stomach in the rat, mouse and guinea-pig. Neuroscience, 25(1):181–93. doi: 10.1016/0306-4522(88)90017-6.

30. Hayakawa T, Kuwahara S, Maeda S, Tanaka K, Seki M. (2009) Distribution of vagal CGRP immunoreactive fibers in the lower esophagus and the cardia of the stomach of the rat. J Chem Neuroanat, 38(2):124–9. doi:10.1016/j.jchemneu.2009.04.001.

31. Hibberd TJ, Yew WP, Dodds KN, Xie Z, Travis L, Brookes SJ, Costa M, Hu H, Spencer NJ. (2022) Quantification of CGRP-immunoreactive myenteric neurons in mouse colon. J Comp Neurol, 530(18):3209–3225. doi: 10.1002/cne.25403

32. Holmberg A, Kaim J, Persson A, Jensen J, Wang T, Holmgren S. (2002) Effects of digestive status on the reptilian gut. Comp Biochem Physiol A Mol Integr Physiol, 133(3):499–518. doi: 10.1016/s1095-6433(02)00257-x.

33. Holzer P. (1998) Neural emergency system in the stomach. Gastroenterology, 114(4):823–39. doi: 10.1016/s0016-5085(98)70597-9.

34. Holzer P. (2000) Local microcirculatory reflexes and afferent signaling in response to gastric acid challenge. Gut, 47 Suppl 4(Suppl 4):iv46–8; discussion iv52. doi: 10.1136/gut.47.suppl_4.iv46.

35. Holzer P. (2006) Efferent-like roles of afferent neurons in the gut: Blood flow regulation and tissue protection. Auton Neurosci, 125(1-2):70–5. doi: 10.1016/j.autneu.2006.01.004.

36. Holzer P, Farzi A. (2014) Neuropeptides and the microbiota-gut-brain axis. Adv Exp Med Biol, 817:195–219. doi: 10.1007/978-1-4939-0897-4_9.

37. Holzer P, Guth PH. (1991) Neuropeptide control of rat gastric mucosal blood flow. Increase by calcitonin gene-related peptide and vasoactive intestinal polypeptide, but not substance P and neurokinin A. Circ Res, 68(1):100–5. doi:10.1161/01.res.68.1.100.

38. Holzer P, Lippe II, Raybould HE, Pabst MA, Livingston EH, Amann R, Peskar BM, Peskar BA, Taché Y, Guth PH. (1991) Role of peptidergic sensory neurons in gastric mucosal blood flow and protection. Ann N Y Acad Sci, 632:272–82. doi: 10.1111/j.1749-6632.1991.tb33115.x.

39. Jaffey DM, Chesney L, Powley TL. (2021) Stomach serosal arteries distinguish gastric regions of the rat. J Anat, 239(4):903–912. doi:10.1111/joa.13480.

40. Jensen J. (1997) Co-release of substance P and neurokinin A from the Atlantic cod stomach. Peptides, 18(5):717–22. doi: 10.1016/s0196-9781(97)00133-2.

41. Julius D, Basbaum AI. (2001) Molecular mechanisms of nociception. Nature, 413(6852):203–10. doi: 10.1038/35093019.

42. Kaestner CL, Smith EH, Peirce SG, Hoover DB. (2019) Immunohistochemical analysis of the mouse celiac ganglion: An integrative relay station of the peripheral nervous system. J Comp Neurol, 527(16):2742–2760. doi: 10.1002/cne.24705.

43. Katsoulis S, Conlon JM. (1989) Calcitonin gene-related peptides relax guinea pig and rat gastric smooth muscle. Eur J Pharmacol, 162(1):129–34. doi:10.1016/0014-2999(89)90612-2.

44. Kee Z, Kodji X, Brain SD. (2018) The Role of Calcitonin Gene Related Peptide (CGRP) in Neurogenic Vasodilation and Its Cardioprotective Effects. *Frontiers in Physiology*, volume 9. doi:10.3389/fphys.2018.01249.

45. Kraenzlin ME, Ch’ng JLC, Mulderry PK, Ghatei MA, Bloom SR. (1985) Infusion of a novel peptide, calcitonin gene-related peptide (CGRP) in man. Pharmacokinetics and effects on gastric acid secretion and on gastrointestinal hormones. Regulatory Peptides, 10(2– 3):189–197, doi.org/10.1016/0167-0115(85)90013-8.

46. Kudrow D, Nguyen L, Semler J, Stroud C, Samaan K, Hoban DB, Wietecha L, Hsu HA, Pearlman E. (2022) A phase IV clinical trial of gastrointestinal motility in adult patients with migraine before and after initiation of a calcitonin gene-related peptide ligand (galcanezumab) or receptor (erenumab) antagonist. Headache, doi: 10.1111/head.14390. Epub ahead of print.

47. Ladic LA, Buchan AM. (1998) Three-dimensional spatial relationship of neuropeptides and receptors in the rat dorsal vagal complex. Brain Res, 795(1-2):312–24. doi: 10.1016/s0006-8993(98)00299-6.

48. Lee SE, Kim DH, Son SM, Choi SY, You RY, Kim CH, Choi W, Kim HS, Lim YJ, Han JY, Kim HW, Yang IJ, Xu WX, Lee SJ, Kim YC, Yun HY. (2020) Physiological function and molecular composition of ATP-sensitive K+ channels in human gastric smooth muscle. J Smooth Muscle Res, 56(0):29–45. doi:10.1540/jsmr.56.29.

49. Lee Y, Shiotani Y, Hayashi N, Kamada T, Hillyard CJ, Girgis SI, MacIntyre I, Tohyama M. (1987) Distribution and origin of calcitonin gene-related peptide in the rat stomach and duodenum: an immunocytochemical analysis. J Neural Transm, 68(1-2):1–14. doi: 10.1007/BF01244635.

50. Lennerz JK, Rühle V, Ceppa EP, Neuhuber WL, Bunnett NW, Grady EF, Messlinger K. (2008) Calcitonin receptor-like receptor (CLR), receptor activity-modifying protein 1 (RAMP1), and calcitonin gene-related peptide (CGRP) immunoreactivity in the rat trigeminovascular system: differences between peripheral and central CGRP receptor distribution. J Comp Neurol, 507(3):1277–99. doi: 10.1002/cne.21607.

51. Leung C, Robbins S, Moss A, Heal M, Osanlouy M, Christie R, Farahani N, Monteith C, Chen J, Hunter P, Tappan S, Vadigepalli R, Cheng ZJ, Schwaber JS (2021) 3D single cell scale anatomical map of sex-dependent variability of the rat intrinsic cardiac nervous system. iScience, 24(7):102795. doi: 10.1016/j.isci.2021.102795.

52. Li L, Hatcher JT, Hoover DB, Gu H, Wurster RD, Cheng ZJ. (2014) Distribution and morphology of calcitonin gene-related peptide and substance P immunoreactive axons in the whole-mount atria of mice. Auton Neurosci, 181:37–48. doi:10.1016/j.autneu.2013.12.010.

53. Luo XJ, Liu B, Dai Z et al. (2013) Stimulation of calcitonin gene-related peptide release through targeting capsaicin receptor: A potential strategy for gastric mucosal protection. Dig Dis Sci 58, 320–325. doi:10.1007/s10620-012-2362-6.

54. Ma J, Mistareehi A, Madas J, Kwiat AM, Bendowski K, Nguyen D, Chen J, Li DP, B Furness J, L Powley T, Cheng ZJ. (2023) Topographical organization and morphology of substance P (SP)-immunoreactive axons in the whole stomach of mice. J Comp Neurol, 531(2):188–216. doi: 10.1002/cne.25386.

55. Maake C, Kloas W, Szendefi M, Reinecke M. (1999) Neurohormonal peptides, serotonin, and nitric oxide synthase in the enteric nervous system and endocrine cells of the gastrointestinal tract of neotenic and thyroid hormone-treated axolotls (Ambystoma mexicanum). Cell Tissue Res, 297(1):91–101. doi: 10.1007/s004410051336.

56. Maggi CA. (1995) Tachykinins and calcitonin gene-related peptide (CGRP) as co-transmitters released from peripheral endings of sensory nerves. Prog Neurobiol, 45(1):1–98. doi:10.1016/0301-0082(94)e0017-b.

57. Makowska K, Gonkowski S. (2019) Age and Sex-Dependent Differences in the Neurochemical Characterization of Calcitonin Gene-Related Peptide-Like Immunoreactive (CGRP-LI) Nervous Structures in the Porcine Descending Colon. Int J Mol Sci, 20(5):1024. doi: 10.3390/ijms20051024.

58. Maton PN, Sutliff VE, Zhou ZC, Collins SM, Gardner JD, Jensen RT. (1988) Characterization of receptors for calcitonin gene-related peptide on gastric smooth muscle cells. Am J Physiol, 254(6 Pt 1):G789–94. doi: 10.1152/ajpgi.1988.254.6.G789.

59. Matsumoto K, Hosoya T, Ishikawa E, Tashima K, Amagase K, Kato S, Murayama T, Horie S. (2014) Distribution of transient receptor potential cation channel subfamily V member 1 expressing nerve fibers in mouse esophagus. Histochem Cell Biol, 142(6):635–44. doi: 10.1007/s00418-014-1246-6.

60. Mawe, G. M., Schemann, M., Wood, J. D., & Gershon, M. D. (1989) Immunocytochemical analysis of potential neurotransmitters present in the myenteric plexus and muscular layers of the corpus of the guinea pig stomach. The Anatomical Record, 224(3), 431–442. doi:10.1002/ar.1092240312.

61. Mazzia C, Clerc N. (1997) Ultrastructural relationships of spinal primary afferent fibres with neuronal and non-neuronal cells in the myenteric plexus of the cat oesophago-gastric junction. Neuroscience, 80(3):925–37. doi: 10.1016/s0306-4522(97)00058-4.

62. Mazzoni M, Clavenzani P, Minieri L, Russo D, Chiocchetti R, Lalatta-Costerbosa G. (2010) Extrinsic afferents supplying the ovine duodenum and ileum. Res Vet Sci, 88(3):361–8. doi: 10.1016/j.rvsc.2009.11.012.

63. McGovern AE, Mazzone SB. (2010) Characterization of the vagal motor neurons projecting to the Guinea pig airways and esophagus. Front Neurol, 1:153. doi: 10.3389/fneur.2010.00153.

64. Meng J, Ovsepian SV, Wang J, Pickering M, Sasse A, Aoki KR, Lawrence GW, Dolly JO. (2009) Activation of TRPV1 mediates calcitonin gene-related peptide release, which excites trigeminal sensory neurons and is attenuated by a retargeted botulinum toxin with anti-nociceptive potential. J Neurosci, 29(15):4981–92. doi: 10.1523/JNEUROSCI.5490-08.2009.

65. Mitsui R. (2009) Characterisation of calcitonin gene-related peptide-immunoreactive neurons in the myenteric plexus of rat colon. Cell Tissue Res,337(1):37–43. doi: 10.1007/s00441-009-0798-6.

66. Mongardi Fantaguzzi C, Thacker M, Chiocchetti R, Furness JB. (2009) Identification of neuron types in the submucosal ganglia of the mouse ileum. Cell Tissue Res, 336(2):179–89. doi: 10.1007/s00441-009-0773-2.

67. Mózsik G, Peidl Z, Szolcsányi J, Dömötör A, Hideg K, Szekeres G, Karádi O, Hunyady B. (2005) Participation of vanilloid/capsaicin receptors, calcitonin-gene-related peptide and substance P in gastric protection of omeprazole and omeprazole-like compounds. Inflammopharmacology, 13(1-3):139–59. doi: 10.1163/156856005774423764.

68. Osanlouy M, Bandrowski A, de Bono B, Brooks D, Cassarà AM, Christie R, Ebrahimi N, Gillespie T, Grethe JS, Guercio LA, Heal M, Lin M, Kuster N, Martone ME, Neufeld E, Nickerson DP, Soltani EG, Tappan S, Wagenaar JB, Zhuang K, Hunter PJ. (2021) The SPARC DRC: Building a Resource for the Autonomic Nervous System Community. Front Physiol,12:693735. doi: 10.3389/fphys.2021.693735.

69. Ouyang A, Broussard DL, Feng HS. (1998) Action of substance P and interaction of calcitonin gene-related peptide and substance P on the cat antral circular muscle. Regul Pept, 77(1-3):25–32. doi: 10.1016/s0167-0115(98)00033-0.

70. Palus K, Bulc M, Całka J. (2018) Changes in VIP-, SP- and CGRP- like immunoreactivity in intramural neurons within the pig stomach following supplementation with low and high doses of acrylamide. Neurotoxicology, 69:47–59. doi: 10.1016/j.neuro.2018.09.002.

71. Parkman HP, Reynolds JC, Elfman KS, Ogorek CP. (1989) Calcitonin gene-related peptide: a sensory and motor neurotransmitter in the feline lower esophageal sphincter. Regul Pept, 25(1):131–46. doi: 10.1016/0167-0115(89)90255-3.

72. Peskar BM, Ehrlich K, Peskar BA. (2002) Role of ATP-sensitive potassium channels in prostaglandin-mediated gastroprotection in the rat. J Pharmacol Exp Ther, 301(3):969–74. doi:10.1124/jpet.301.3.969.

73. Poonyachoti S, Kulkarni-Narla A, Brown DR. (2002) Chemical coding of neurons expressing delta- and kappa-opioid receptor and type I vanilloid receptor immunoreactivities in the porcine ileum. Cell Tissue Res, 307(1):23–33. doi: 10.1007/s00441-001-0480-0.

74. Powley TL. (2021) Brain-gut communication: vagovagal reflexes interconnect the two“brains”. Am J Physiol Gastrointest Liver Physiol, 321: G576–G587. doi:10.1152/ajpgi.00214.2021.

75. Powley TL, Hudson CN, McAdams JL, Baronowsky EA, Phillips RJ. (2016) Vagal Intramuscular Arrays: The Specialized Mechanoreceptor Arbors That Innervate the Smooth Muscle Layers of the Stomach Examined in the Rat. J Comp Neurol, 524(4):713–37. doi: 10.1002/cne.23892.

76. Powley TL, Jaffey DM, McAdams J, Baronowsky EA, Black D, Chesney L, Evans C, Phillips RJ. (2019) Vagal innervation of the stomach reassessed: brain-gut connectome uses smart terminals. Ann N Y Acad Sci, 1454(1):14–30. doi: 10.1111/nyas.14138.

77. Qu ZD, Thacker M, Castelucci P, Bagyánszki M, Epstein ML, Furness JB. (2008) Immunohistochemical analysis of neuron types in the mouse small intestine. Cell Tissue Res, 334(2):147–61. doi: 10.1007/s00441-008-0684-7.

78. Ren J, Gao J, Ojeas H, Lightfoot SA, Kida M, Brewer K, Harty RF. (2000) Involvement of capsaicin-sensitive sensory neurons in stress-induced gastroduodenal mucosal injury in rats. Dig Dis Sci, 45(4):830–6. doi: 10.1023/a:1005424617101.

79. Rodrigo J, Polak JM, Fernandez L, Ghatei MA, Mulderry P, Bloom SR. (1985) Calcitonin gene related peptide immunoreactive sensory and motor nerves of the rat, cat, and monkey esophagus. Gastroenterology, 88(2):444–51. doi: 10.1016/0016-5085(85)90505-0.

80. Rosenfeld M, Mermod JJ, Amara SG, Swanson LW, Sawchenko PE, Rivier J, Vale WW, Evans RM. (1983) Production of a novel neuropeptide encoded by the calcitonin gene via tissue-specific RNA processing. Nature 304:129–135. doi:10.1038/304129a0.

81. Rudolf K, Eberlein W, Engel W, Pieper H, Entzeroth M, Hallermayer G, Doods H. (2005) Development of human calcitonin gene-related peptide (CGRP) receptor antagonists. 1. Potent and selective small molecule CGRP antagonists. 1-[N2-[3,5-dibromo-N-[[4-(3,4-dihydro-2(1H)-oxoquinazolin-3-yl)-1-piperidinyl]carbonyl]-D-tyrosyl]-l-lysyl]-4-(4-pyridinyl)piperazine: the first CGRP antagonist for clinical trials in acute migraine. J Med Chem, 48(19):5921–31. doi: 10.1021/jm0490641.

82. Russell FA, King R, Smillie SJ, Kodji X, Brain SD. (2014) Calcitonin gene-related peptide: physiology and pathophysiology. Physiol Rev, 94(4):1099–142. doi:10.1152/physrev.00034.2013.

83. Rysevaite K, Saburkina I, Pauziene N, Vaitkevicius R, Noujaim SF, Jalife J, Pauza DH. (2011) Immunohistochemical characterization of the intrinsic cardiac neural plexus in whole mount mouse heart preparations. Heart Rhythm, 8(5):731–8. doi: 10.1016/j.hrthm.2011.01.013.

84. Rytel L, Palus K, Całka J. (2015) Co-expression of PACAP with VIP, SP and CGRP in the porcine nodose ganglion sensory neurons. Anat Histol Embryol, 44(2):86–91. doi: 10.1111/ahe.12111.

85. Rytel L, Całka J. (2016) Neuropeptide profile changes in sensory neurones after partial prepyloric resection in pigs. Ann Anat, 206:48–56. doi: 10.1016/j.aanat.2016.03.003.

86. Serzysko T, Skwarek A, Chudziak E, Malina M, Kaleczyc J, Sienkiewicz W. (2021) Enteric neuronal development in canine small intestine - an immunohistochemical study. Pol J Vet Sci, 24(2):293–301. doi: 10.24425/pjvs.2021.137665.

87. Sharrad DF, Hibberd TJ, Kyloh MA, Brookes SJ, Spencer NJ. (2015) Quantitative immunohistochemical co-localization of TRPV1 and CGRP in varicose axons of the murine oesophagus, stomach, and colorectum. Neurosci Lett, 599:164–71. doi:10.1016/j.neulet.2015.05.020.

88. Shochina M, Belai A, Toole L, Knight G, Burnstock G. (1997) Neurochemical coding in the myenteric plexus of the upper gastrointestinal tract of hibernating hamsters. Int J Dev Neurosci, 15(3):353–62. doi: 10.1016/s0736-5748(97)00003-8.

89. Smolilo DJ, Hibberd TJ, Costa M, Wattchow DA, De Fontgalland D, Spencer NJ. (2020) Intrinsic sensory neurons provide direct input to motor neurons and interneurons in mouse distal colon via varicose baskets. J Comp Neurol, 528(12):2033–2043. doi: 10.1002/cne.24872.

90. Spencer NJ, Greenheigh S, Kyloh M, Hibberd TJ, Sharma H, Grundy L, Brierley SM, Harrington AM, Beckett EA, Brookes SJ, Zagorodnyuk VP. (2018) Identifying unique subtypes of spinal afferent nerve endings within the urinary bladder of mice. J Comp Neurol, 526(4):707–720. doi: 10.1002/cne.24362.

91. Spencer NJ, Kyloh M, Beckett EA, Brookes S, Hibberd T. (2016) Different types of spinal afferent nerve endings in stomach and esophagus identified by anterograde tracing from dorsal root ganglia. J Comp Neurol, 524(15):3064–83. doi: 10.1002/cne.24006.

92. Spencer NJ, Kyloh M, Duffield M. (2014) Identification of Different Types of Spinal Afferent Nerve Endings That Encode Noxious and Innocuous Stimuli in the Large Intestine Using a Novel Anterograde Tracing Technique. PLoS ONE, 9(11): e112466. https://doi.org/10.1371/journal.pone.0112466.

93. Spencer NJ, Kyloh MA, Travis L, Dodds KN. (2020) Identification of spinal afferent nerve endings in the colonic mucosa and submucosa that communicate directly with the spinal cord: The gut-brain axis. J Comp Neurol, 528(10):1742–1753. doi: 10.1002/cne.24854.

94. Sternini C, Anderson K. (1992) Calcitonin Gene-Related Peptide-Containing Neurons Supplying the Rat Digestive System: Differential Distribution and Expression Pattern. Somatosensory & Motor Research, 9(1), 45–59. doi:10.3109/08990229209144762

95. Sternini C, De Giorgio R, Furness JB. (1992) Calcitonin gene-related peptide neurons innervating the canine digestive system. Regul Pept, 42(1-2):15–26. doi:10.1016/0167-0115(92)90020-u.

96. Sternini C, Reeve JR Jr, Brecha N. (1987) Distribution and characterization of calcitonin gene related peptide immunoreactivity in the digestive system of normal and capsaicin-treated rats. Gastroenterology, 93(4):852–62. doi: 10.1016/0016-5085(87)90450-1

97. Su HC, Bishop AE, Power RF, Hamada Y, Polak JM. (1987) Dual intrinsic and extrinsic origins of CGRP- and NPY-immunoreactive nerves of rat gut and pancreas. J Neurosci, 7(9):2674–87. doi: 10.1523/JNEUROSCI.07-09-02674.1987.

98. Sun T, Guo Z, Liu CJ, Li MR, Li TP, Wang X, Yuan DJ. (2016) Preservation of CGRP in myocardium attenuates development of cardiac dysfunction in diabetic rats. Int J Cardiol, 220:226–34. doi: 10.1016/j.ijcard.2016.06.092.

99. Suzuki T, Kagoshima M, Shibata M, Inaba N, Onodera S, Yamaura T, Shimada H. (1997) Effects of several denervation procedures on distribution of calcitonin gene-related peptide and substance P immunoreactive in rat stomach. Dig Dis Sci, 42(6):1242–54. doi: 10.1023/a:1018858208532.

100. Szolcsányi J. (2004) Forty years in capsaicin research for sensory pharmacology and physiology. Neuropeptides, 38(6):377–84. doi: 10.1016/j.npep.2004.07.005.

101. Tan LL, Bornstein JC, Anderson CR. (2008) Distinct chemical classes of medium-sized transient receptor potential channel vanilloid 1-immunoreactive dorsal root ganglion neurons innervate the adult mouse jejunum and colon. Neuroscience, 156(2):334–43. doi: 10.1016/j.neuroscience.2008.06.071.

102. Tarif AMM, Islam MN, Jahan MR, Yanai A, Nozaki K, Masumoto KH, Shinoda K. (2021) Immunohistochemical expression and neurochemical phenotypes of huntingtin-associated protein 1 in the myenteric plexus of mouse gastrointestinal tract. Cell Tissue Res, 386(3):533–558. doi: 10.1007/s00441-021-03542-4.

103. Thomson M. (2017) Embryology of the Stomach. Springer-Verlag: Esophageal and Gastric Disorders in Infancy and Childhood, (110)1253–61 doi:10.1007/978-3-642-11202-7_110.

104. Timmermans JP, Scheuermann DW, Barbiers M, Adriaensen D, Stach W, Van Hee R, De Groodt-Lasseel MH. (1992) Calcitonin gene-related peptide-like immunoreactivity in the human small intestine. Acta Anat (Basel), 143(1):48–53. doi:10.1159/000147227.

105. Timmermans JP, Adriaensen D, Cornelissen W, Scheuermann DW. (1997) Structural organization and neuropeptide distribution in the mammalian enteric nervous system, with special attention to those components involved in mucosal reflexes. Comparative Biochemistry and Physiology Part A: Physiology, 118(2), 331–340. doi: 10.1016/S0300-9629(96)00314-3.

106. Toole L, Belai A, Shochina M, Burnstock G. (1999) The effects of hibernation on the myenteric plexus of the golden hamster small and large intestine. Cell Tissue Res, 296(3):479–87. doi: 10.1007/s004410051308.

107. van Rossum D, Hanisch UK, Quirion R. (1997) Neuroanatomical localization, pharmacological characterization and functions of CGRP, related peptides and their receptors. Neurosci Biobehav Rev, 21(5):649–78. doi:10.1016/s0149-7634(96)00023-1.

108. Walter GC, Phillips RJ, McAdams JL, Powley TL. (2016) Individual sympathetic postganglionic neurons co-innervate myenteric ganglia and smooth muscle layers in the gastrointestinal tract of the rat. J Comp Neurol, 524(13):2577–603. doi: 10.1002/cne.23978.

109. Wang FB, Powley TL. (2000) Topographic inventories of vagal afferents in gastrointestinal muscle, J Comp Neurol, 421: 302–324. https://doi.org/10.1002/(SICI)1096-9861(20000605)421:3<302::AID-CNE2>3.0.CO;2-N.

110. Warzecha Z, Dembiński A, Ceranowicz P, Stachura J, Tomaszewska R, Konturek SJ. (2001) Effect of sensory nerves and CGRP on the development of caerulein-induced pancreatitis and pancreatic recovery. J Physiol Pharmacol, 52(4 Pt 1):679–704.

111. Yaprak M. (2008) The axon reflex. Neuroanatomy, 7:17–19.

112. Zhong F, Christianson JA, Davis BM, Bielefeldt K. (2008) Dichotomizing axons in spinal and vagal afferents of the mouse stomach. Dig Dis Sci, 53(1):194–203. doi: 10.1007/s10620-007-9843-z.

